# Local translation couples synaptic activity to mitochondrial adaptation in dendrites

**DOI:** 10.64898/2026.09.17.752159

**Authors:** Madison T. Jones, Natalie Noble, Robert B. Darnell, Ezgi Hacisuleyman

## Abstract

Neurons rely on localized protein synthesis to rapidly adapt synaptic function to activity, yet how dendritic translation regulates mitochondrial remodeling during synaptic plasticity remains poorly understood. Here, we show that neuronal activity engages a spatially restricted translational program that couples local protein synthesis to mitochondrial function through the non-canonical translation initiation factor eIF4G2. Using proximity labeling to profile the dendritic RNA interactome, translatome, and proteome, we identify a cohort of nuclear-encoded mitochondrial mRNAs that are selectively recruited for translation following depolarization and mGluR activation. This program drives activity-dependent increases in mitochondrial membrane potential, mitochondrial abundance, and oxygen consumption. Loss of eIF4G2 abolishes these responses, whereas dendrite-specific, but not soma-restricted, rescue restores mitochondrial remodeling, demonstrating that eIF4G2 functions locally at postsynaptic sites. Mechanistically, eIF4G2 binds the 5′ untranslated regions (5’UTRs) of activity-responsive mitochondrial transcripts and promotes translation of both upstream open reading frames (uORFs) and downstream coding sequences. Using a dendritically targeted split-GFP reporter, we further show that neuronal activity induces local uORF translation to generate previously unannotated micropeptides. Together, our findings identify eIF4G2-dependent local translation as a mechanism that establishes mitochondrial competence during synaptic activity by coordinating the production of mitochondrial proteins and uORF-encoded micropeptides.

**One sentence summary:** Neuronal activity engages dendritic eIF4G2-dependent 5′UTR translation of mitochondrial proteins and micropeptides to remodel local mitochondria. (145 characters)

## Introduction

Synaptic activity imposes rapid and spatially restricted energetic and metabolic demands that must be met locally within dendrites^1^. Mitochondria are central to this adaptation by generating ATP, buffering intracellular calcium, and supplying metabolites required for neurotransmission, protein synthesis, and structural remodeling of synapses^2^. Rather than functioning as static energy producers, mitochondria undergo activity-dependent changes in positioning, morphology, membrane potential, and respiratory function^3,4^. These responses have been studied most extensively at presynaptic terminals, where they support neurotransmitter release and synaptic transmission^5,6^. By contrast, the mechanisms that dynamically regulate mitochondrial function within dendrites and postsynaptic compartments remain far less well understood, despite the positioning of dendritic mitochondria to meet the local energetic and metabolic demands imposed by synaptic plasticity. How individual synapses rapidly adapt mitochondrial function, and how synaptic signaling is coupled to localized regulation of mitochondrial proteomes and physiology, therefore remain fundamental unanswered questions.

Neurons overcome the constraints imposed by their elaborate morphology through local translation, allowing proteins to be synthesized directly within dendrites rather than transported over long distances from the soma^7–11^. Following neuronal activation, hundreds of dendritically localized mRNAs undergo rapid translational regulation^12^, enabling compartment-specific remodeling of receptors, signaling molecules, cytoskeletal proteins, and other factors required for synaptic plasticity^13–18^. While considerable progress has been made in identifying dendritic transcripts and the RNA-binding proteins^17,19^ that regulate their localization, transport, and translation, much less is known about the physiological consequences of activity-dependent local translation. In particular, whether dendritic protein synthesis directly remodels mitochondrial composition and function to meet the energetic demands of active synapses remains largely unexplored.

Although mitochondria possess their own genome, more than 99% of mitochondrial proteins are encoded by nuclear genes, translated on cytosolic ribosomes, and subsequently imported into the organelle. This organization raises the intriguing possibility that localized translation of nuclear-encoded mitochondrial transcripts could provide an efficient mechanism for rapidly adapting mitochondrial function at individual synapses. Indeed, numerous mitochondrial mRNAs have been detected within neuronal processes^12,13^, suggesting that neurons locally position the molecular components required for mitochondrial remodeling. However, whether neuronal activity selectively regulates the translation of these transcripts, how this process is spatially restricted within dendrites, and whether locally synthesized mitochondrial proteins directly contribute to mitochondrial adaptation have remained unresolved.

Progress toward addressing these questions has been limited by the difficulty of interrogating localized RNA-protein interactions and translation within intact neuronal compartments. Early imaging and biochemical studies established the presence of ribosomes and mRNAs within dendrites^20–22^, whereas transcriptomic approaches identified thousands of localized RNAs together with the cis-elements and RNA-binding proteins governing their localization and stability^13,15,23–29^. More recently, proximity-labeling technologies have enabled high-resolution mapping of subcellular transcriptomes and proteomes in living cells^30–32^. However, approaches capable of simultaneously capturing localized RNA binding, translation, and protein production during rapid neuronal activation have remained limited.

We previously developed a PSD95-targeted proximity-labeling platform coupled to crosslinking immunoprecipitation (PL-CLIP), ribosome profiling (PL-Ribo-seq) and quantitative mass spectrometry (PL-MS) to interrogate RNA localization, local translation, and protein abundance within postsynaptic dendrites^12^. Using this platform, we showed that neuronal activity rapidly reprograms dendritic translation and identified the non-canonical translation initiation factor eIF4G2 and upstream open reading frame (uORF)-containing transcripts as central components of this response. These findings raised the possibility that eIF4G2-dependent translation coordinates broader functional adaptations within activated dendrites, but its compartment-specific RNA interactions and physiological consequences remained unknown.

uORFs are increasingly recognized as dynamic regulators of gene expression that can influence translation of downstream coding sequences^33,34^. Ribosome-profiling studies have further revealed pervasive translation of uORFs and other noncanonical short open reading frames, some of which encode functional micropeptides with diverse cellular roles^35–37^. More broadly, translation of such previously unrecognized ORFs may provide a substrate for evolutionary innovation, enabling initially noncanonical peptides to acquire biological functions and, in some cases, contribute to de novo gene emergence^38–40^. However, whether neuronal activity dynamically induces uORF translation within dendrites, whether these events generate previously unannotated local micropeptides, and how uORF and coding-sequence translation are coordinated at active synapses remain unknown. It is also unclear whether a localized translational program can couple the production of activity-regulated micropeptides with remodeling of the dendritic mitochondrial proteome.

Here, we further develop the CLIP arm of this platform to map activity-dependent eIF4G2-RNA interactions at nucleotide resolution specifically within the postsynaptic compartment. We integrate this expanded PL-CLIP workflow with PL-Ribo-seq, PL-MS, live-cell imaging, and functional analyses of mitochondrial physiology to determine how eIF4G2-dependent local translation remodels dendritic mitochondria during synaptic activation. PL-CLIP reveals activity-dependent eIF4G2 binding to the 5’ untranslated regions (5′UTRs) of mitochondrial transcripts and links these interactions to the translation of both uORFs and downstream coding sequences. We identify a translational program that promotes the local synthesis of nuclear-encoded mitochondrial proteins, driving activity-dependent increases in mitochondrial membrane potential, mitochondrial mass, and oxidative metabolism. Finally, we show that neuronal activity induces local uORF translation to generate previously unannotated dendritic micropeptides, uncovering an unexpected layer of activity-dependent gene regulation. Together, our findings demonstrate the expanded utility of PL-CLIP for resolving compartment-specific RNA-protein interactions and identify eIF4G2-dependent local translation as a mechanism that establishes mitochondrial competence at active synapses through the coordinated production of mitochondrial proteins and uORF-encoded micropeptides.

## Results

### Activity-dependent dendritic translation preferentially targets nuclear-encoded mitochondrial proteins

To determine how activity-dependent changes in dendritic RNA regulation are propagated through translation to the local proteome, we integrated postsynaptic PL-CLIP, PL-Ribo-seq, and PL-MS datasets from resting and activated neurons^12^. Primary cortical neurons were analyzed under resting conditions or following KCl-mediated depolarization (hereafter, Dep) or stimulation with the group I mGluR agonist DHPG, allowing activity-dependent changes in RNA binding, translation, and protein abundance to be compared simultaneously.

Neuronal activation remodeled multiple functional classes in the dendritic translatome and proteome, including synaptic signaling, cytoskeletal organization, RNA regulation, and protein modification. However, nuclear-encoded mitochondrial transcripts and proteins emerged as the most dominant enriched functional class following both KCl and DHPG stimulation (Fig. 1a,b and Extended Data Fig. 1a-d). Their enrichment in PL-Ribo-seq and PL-MS was not accompanied by corresponding changes in their local RNA abundance (Fig. 1c), indicating regulation primarily at the level of translation rather than transcript localization or stability. Although KCl induced broader translational changes across multiple functional categories, DHPG preferentially promoted translation of mitochondrial transcripts and 3′UTR-regulated mRNAs, in several cases preceding detectable changes in steady-state protein abundance (Fig. 1a,b and Extended Data Fig. 1c,d). Despite these stimulus-specific differences, both activation paradigms converged on enhanced local translation of nuclear-encoded mitochondrial proteins, identifying mitochondrial function as a prominent feature of activity-dependent dendritic remodeling.

**Fig. 1.**
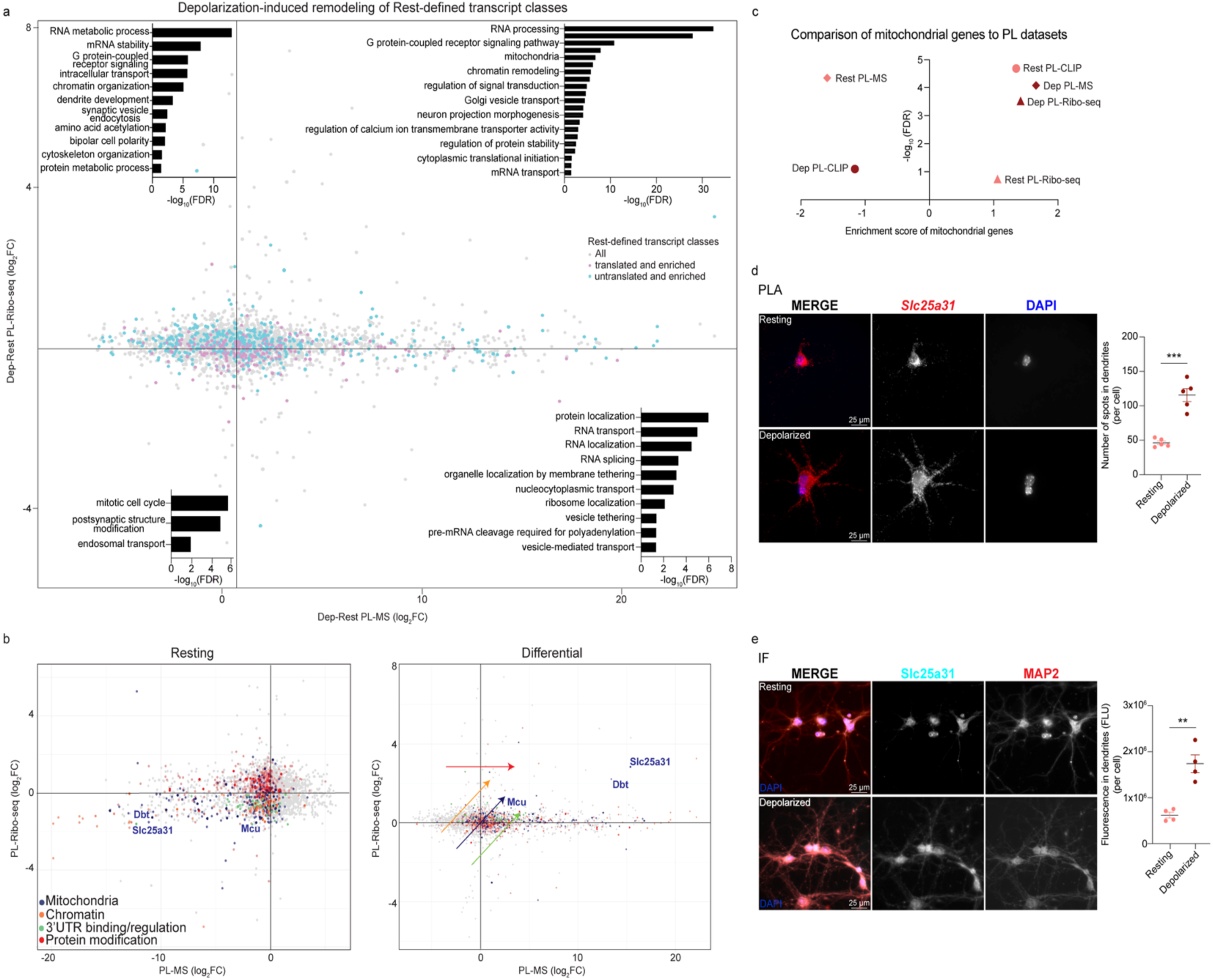
Synaptic activation elevates mitochondrial mRNA translation and reshapes dendritic RNA programs. **a**, Gene Ontology (GO) enrichment analysis of activity-dependent (Dep-Rest) changes measured by PL-MS (n= 5 biological replicates) and PL-Ribo-seq (n= 3 biological replicates). GO terms were derived from Dep-Rest differential analyses. Colored distributions denote transcript classes defined in the resting-state dataset, with purple indicating dendritically enriched and translated transcripts and cyan indicating dendritically enriched but untranslated transcripts. Significantly enriched terms are shown at FDR < 0.05 following Benjamini–Hochberg correction. **b**, Distribution of mitochondria, chromatin, 3’UTR regulation, and protein modification-related RNAs shown in resting and differential PL-MS vs. PL-Ribo-seq. **c**, Dendritic mitochondrial RNAs compared to ranked PL-CLIP, PL-Ribo-seq, and PL-MS datasets from resting and depolarized neurons. **d,e,** Nascent *Slc25a31* protein synthesis (**d**) and total SLC25A31 protein levels (**e**) in dendrites were measured using puromycin proximity ligation assay (puro-PLA; n= 5 biological replicates, p_adj_= 1.0x10^-4^) and immunofluorescence (IF; n= 4 biological replicates, p_adj_= 1.6x10^-3^), respectively. For each biological replicate, three fields containing 25-35 neurons per field were analyzed, and the field-level measurements were averaged to obtain one value per biological replicate. DAPI labels nuclei, and MAP2 demarcates dendrites. All data are presented as mean ± SEM. Statistical significance was determined by one-way ANOVA followed by Tukey’s multiple comparisons test. Scale bars= 25 μm.

To validate these sequencing-and proteomics-based observations, we measured nascent synthesis of the nuclear-encoded mitochondrial protein SLC25A31 using puromycin-based proximity ligation assays (puro-PLA) and quantified endogenous SLC25A31 abundance by immunofluorescence (IF). Both approaches revealed increased dendritic signal following KCl depolarization and DHPG stimulation (Fig. 1d,e and Extended Data Fig. 2a,b). In contrast, mitochondrial fission factor (MFF), another nuclear-encoded mitochondrial protein, showed no comparable increase (Extended Data Fig. 2c,d). These results indicate that neuronal activity selectively increases the local synthesis and dendritic abundance of specific mitochondrial proteins rather than producing a general increase in mitochondrial protein abundance.

Together, these findings raised the possibility that activity-dependent local translation serves not only to remodel dendritic protein composition but directly adapts mitochondrial physiology to the energetic demands of synaptic activation. We therefore asked whether this translational program directly regulates mitochondrial function.

### Neuronal activity drives calcium-dependent remodeling of dendritic mitochondria

The preferential translation of nuclear-encoded mitochondrial proteins suggested that activity-dependent local translation may directly remodel mitochondrial physiology within dendrites. To test this possibility, we quantified mitochondrial mass and membrane potential (Δψm) using MitoTracker Green FM and MitoTracker Deep Red FM, respectively. Both KCl depolarization and DHPG stimulation significantly increased dendritic mitochondrial abundance and Δψm, demonstrating that neuronal activity rapidly enhances mitochondrial remodeling and function (Fig. 2a). Because synaptic activity is accompanied by calcium influx, we next asked whether these responses required extracellular calcium. Chelation with EGTA abolished the activity-dependent increases in both mitochondrial mass and membrane potential following either stimulation paradigm, indicating that rapid mitochondrial remodeling within dendrites is calcium dependent (Fig. 2a).

**Fig. 2.**
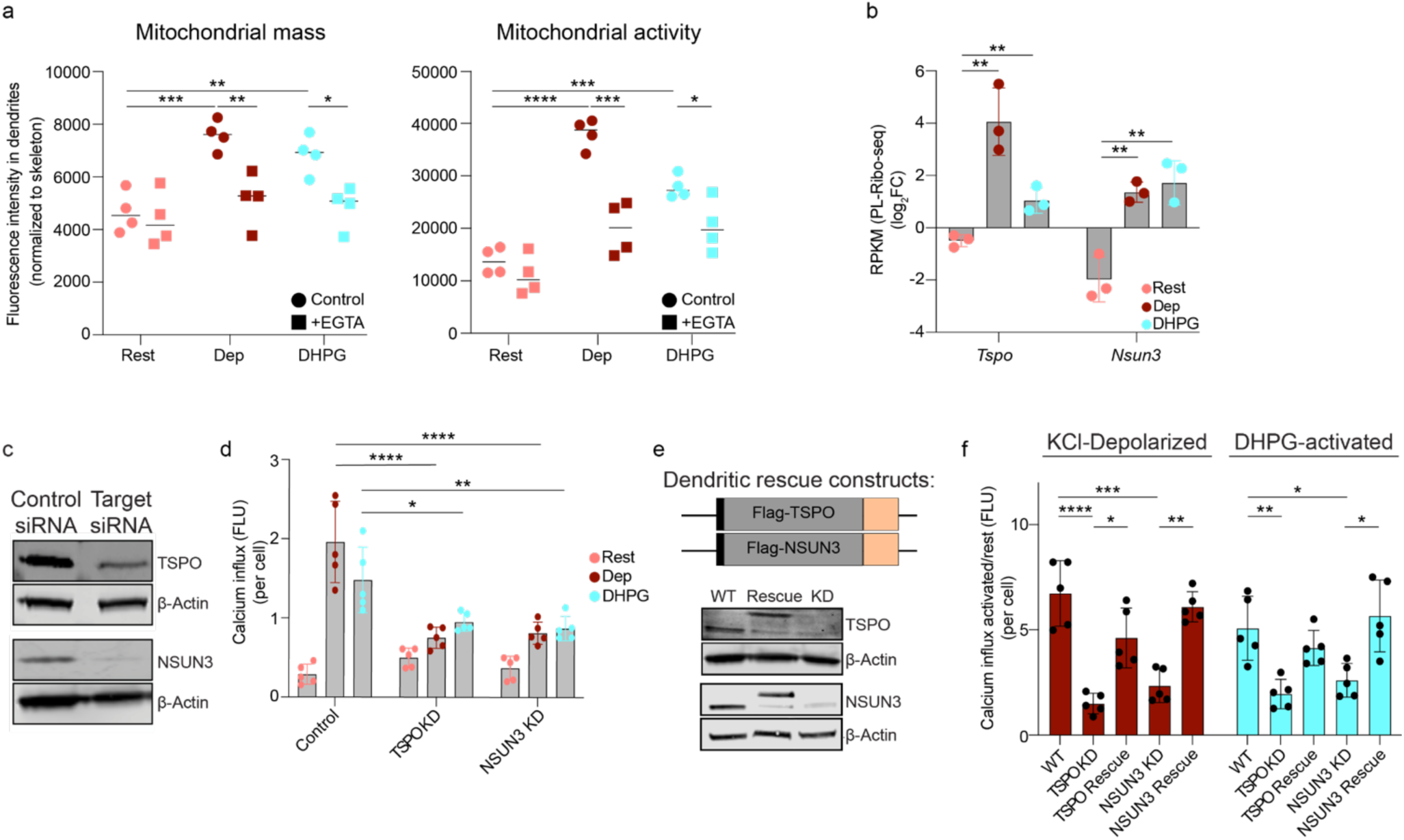
Calcium-dependent mitochondrial activation and dendritic translation of *Tspo* and *Nsun3* support neuronal calcium homeostasis. **a**, Quantification of dendritic mitochondrial mass and membrane potential (Δψm) using Mitotracker Green and Deep Red, respectively, in resting and depolarized (KCl or DHPG-treated) neurons. Median values are shown. Mitochondrial mass: Rest vs. Dep Control p_adj_= 9.2x10^-4^; Dep Control vs. +EGTA p_adj_= 5.7x10^-3^; Rest vs. DHPG Control p_adj_= 6.4x10^-3^; DHPG Control vs. +EGTA p_adj_= 1.0x10^-2^. Mitochondrial activity: Rest vs. Dep Control p_adj_= 1.4x10^-5^; Dep Control vs. +EGTA p_adj_= 8.1x10^-4^; Rest vs. DHPG Control p_adj_= 1.5x10^-4^; DHPG Control vs. +EGTA p_adj_= 3.2x10^-2^. **b**, Dendritic translation of *Tspo* and *Nsun3* measured by PL-Ribo-seq (log_2_, CDS) in resting, KCl-, and DHPG-stimulated neurons (n= 3 biological replicates). Significance is calculated using the two-tailed, unpaired student’s t-test. Data are presented as mean ± SD. **c**, Western blot validation of siRNA knockdown (KD) of TSPO and NSUN3 in whole neurons, shown with β-actin loading control. **d**, Activity-dependent calcium influx measured using Fluo4-AM in control, TSPO KD, and NSUN3 KD neurons (n= 5 biological replicates). Dep (KCl): TSPO KD, p_adj_ = 3.5x10⁻⁵; NSUN3 KD, p_adj_ = 6.4x10⁻⁵ (all compared to control). DHPG: TSPO KD, p_adj_ = 0.017; NSUN3 KD, p_adj_ = 0.0061 (all compared to control). **e**, Schematic of dendrite-targeted rescue constructs and corresponding Western blots validating knockdown and localized re-expression of TSPO and NSUN3. **f**, Calcium influx in KD and rescue conditions following KCl or DHPG stimulation (Fluo-4 AM; n = 5 biological replicates). KCl: WT vs. Tspo KD, p_adj_= 1.9x10^-5^; WT vs. Nsun3 KD, p_adj_= 1.5x10^-4^; Tspo KD vs. Rescue, padj= 1.6x10^-2^; Nsun3 KD vs. Rescue, p_adj_= 3.3x10^-3^. DHPG: WT vs. Tspo KD, p_adj_= 6.4x10^-3^; WT vs. Nsun3 KD, p_adj_= 2.9x10^-2^; Nsun3 KD vs. Rescue, p_adj_= 1.9x10^-2^. **d, f,** Data are presented as mean ± SD. Statistical significance was determined by one-way ANOVA followed by Tukey’s multiple comparisons test.

To identify candidate effectors underlying this activity-dependent mitochondrial remodeling, we next examined individual nuclear-encoded mitochondrial transcripts identified by PL-Ribo-seq. Among the activity-responsive mitochondrial transcripts, *Tspo* and *Nsun3* displayed some of the strongest increases in dendritic translation following both KCl depolarization and DHPG stimulation (Fig. 2b). Because TSPO and NSUN3 have established roles in mitochondrial physiology and mitochondrial RNA metabolism, respectively, we selected them as representative candidate effectors for subsequent mechanistic analyses of activity-dependent mitochondrial adaptation. Their robust translational induction further suggested that neuronal activity coordinately regulates multiple aspects of mitochondrial biology, encompassing both mitochondrial function and maintenance.

The calcium dependence of activity-induced mitochondrial remodeling suggested a close functional relationship between mitochondrial adaptation and calcium signaling. Mitochondria both respond to calcium influx and shape intracellular calcium dynamics. Indeed, glutamate receptor-evoked calcium entry has been reported to regulate Miro1-dependent mitochondrial positioning at active synapses, whereas PDZD8-mediated ER-mitochondria contacts facilitate mitochondrial calcium uptake and shape dendritic calcium dynamics^41,42^. These findings establish a bidirectional relationship between mitochondrial organization and neuronal calcium signaling, yet the molecular mechanisms coupling this relationship to activity-dependent local translation remain poorly understood. Because *Tspo* and *Nsun3* were among the most strongly translated mitochondrial transcripts following neuronal activation (Fig. 2b), we asked whether their local synthesis contributes to activity-dependent calcium responses.

To test this, we depleted TSPO or NSUN3 throughout neurons using siRNAs and monitored calcium responses following neuronal stimulation. As benchmark perturbations of neuronal calcium homeostasis, we also depleted CAMK2α, an established regulator of excitatory synaptic signaling and plasticity^43^, and PMCA2, a plasma membrane calcium ATPase that mediates calcium extrusion (Extended Data Fig. 3). As expected, CAMK2α depletion reduced calcium responses following both KCl depolarization and DHPG stimulation, whereas PMCA2 depletion increased stimulus-evoked calcium levels, demonstrating that our assay detects modulation of neuronal calcium signaling in both directions (Extended Data Fig. 3). Under the same conditions, depletion of either TSPO or NSUN3 significantly reduced activity-dependent calcium responses (Fig. 2c,d). These findings suggest that activity-induced synthesis of mitochondrial proteins in dendrites helps establish the mitochondrial state needed to support subsequent calcium signaling. Because these experiments depleted each protein throughout the neuron, they did not establish whether its dendritic synthesis was important. We therefore re-expressed TSPO or NSUN3 using dendritically targeted rescue constructs (Fig. 2e). Local re-expression of either protein restored the activity-dependent calcium response in its respective knockdown background (Fig. 2f), demonstrating that local translation of individual mitochondrial proteins is sufficient to support synaptic calcium homeostasis. These results suggest that the local translation of mitochondrial proteins helps establish the mitochondrial competence needed to sustain calcium signaling following neuronal activation.

### PL-CLIP identifies activity-dependent eIF4G2 interactions with mitochondrial transcripts in dendrites

Having identified an activity-dependent translational program that promotes local synthesis of nuclear-encoded mitochondrial proteins, we next asked how these transcripts are selectively recognized within dendrites. We previously showed, using whole-cell CLIP and transcriptome-wide analyses, that the non-canonical translation initiation factor eIF4G2 preferentially associates with the 5’UTRs of activity-responsive mRNAs to promote downstream translation^12^. However, these approaches lacked the spatial resolution to distinguish RNA interactions occurring specifically within dendrites.

To address this limitation, we adapted our PL-CLIP strategy to map activity-dependent eIF4G2-RNA interactions specifically within postsynaptic dendrites of TurboID-PSD95-transduced neurons (Fig. 3a,b). Sequential eIF4G2 immunoprecipitation followed by streptavidin enrichment robustly recovered dendritically biotinylated eIF4G2-RNA complexes (Extended Data Fig. 4a,b), demonstrating that PL-CLIP can resolve the compartment-specific RNA interactome of a defined RNA-binding protein.

**Fig. 3.**
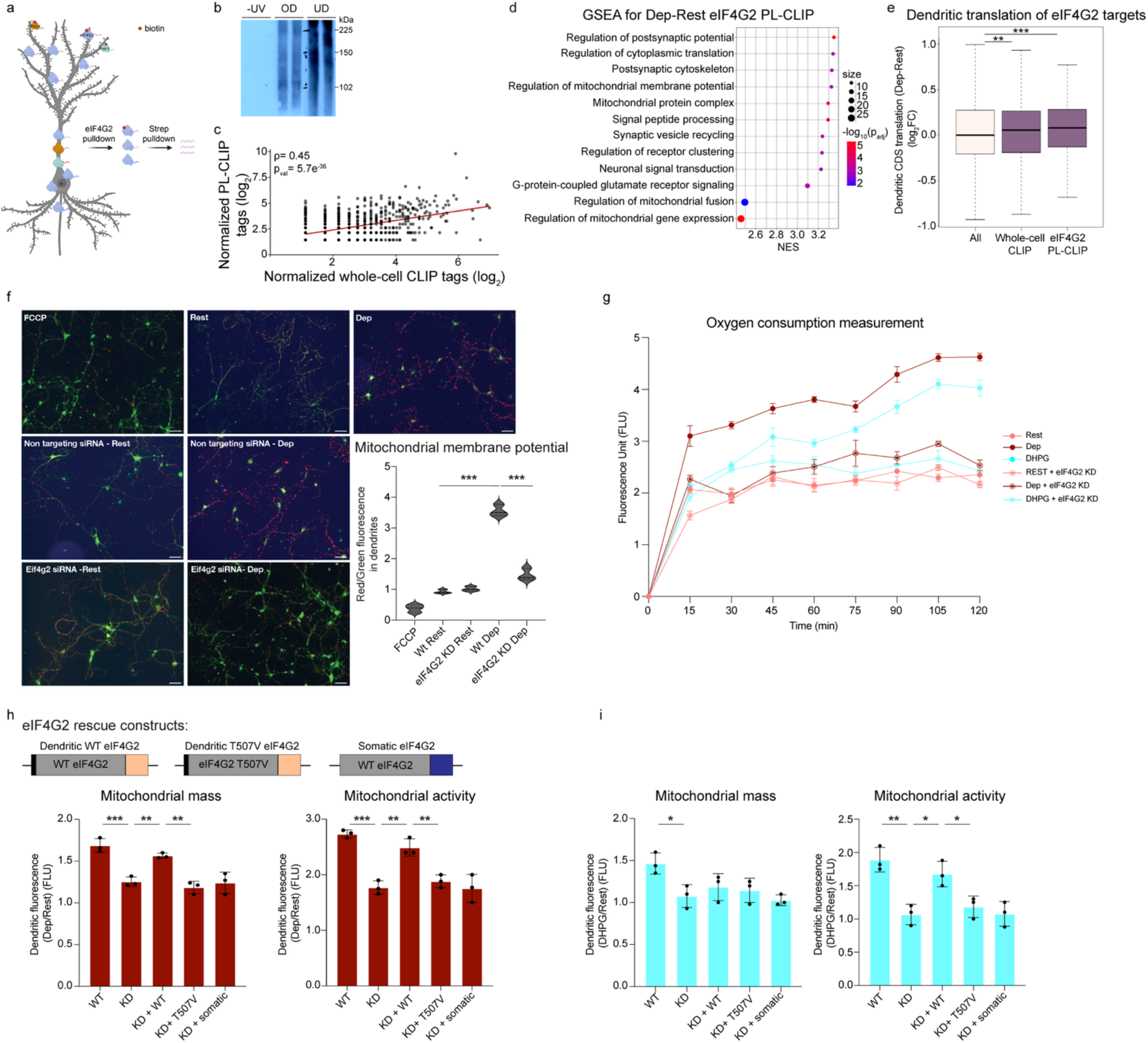
eIF4G2 binds 5′UTRs of dendritic mitochondrial RNAs and regulates local translation and bioenergetics. **a**, Schematic of the eIF4G2 proximity-Labeled CLIP (PL-CLIP) workflow, coupling TurboID-mediated postsynaptic proximity labeling with CLIP-based RNA-protein interaction profiling. **b**, Autoradiograph of eIF4G2 PL-CLIP under -UV control and RNase overdigestion (OD) and underdigestion (UD) conditions. **c**, Correlation between conventional whole-cell eIF4G2 CLIP and dendritic PL-CLIP across transcripts detected by both approaches under depolarizing conditions, demonstrating concordance between total and compartment-resolved eIF4G2 RNA binding. **d**, Gene set enrichment analysis (GSEA) of differentially eIF4G2-bound RNAs identified by PL-CLIP across annotated mitochondrial and synaptic gene sets. Significantly enriched gene sets are shown at FDR < 0.05, with multiple-testing correction performed using the Benjamini-Hochberg method. **e**, Dendritic CDS translation by PL-Ribo-seq (Dep-Rest) for eIF4G2-bound RNAs identified by whole-cell CLIP (p_val_= 3.4x10^-3^) and dendritic PL-CLIP (p_val_= 5.0x10^-4^). Statistical significance was assessed using a two-sided paired Wilcoxon signed-rank test. **f**, JC-1 assay of mitochondrial membrane potential (Δψm) under FCCP control, Rest, and Dep conditions in wild type (WT) and eIF4G2 knockdown (KD) neurons, quantified as the red/green fluorescence ratio. Wt Rest vs. Dep: p_adj_= 6.6x10^-8^ and Wt Dep vs. eIF4G2 KD Dep: p_adj_= 5.5x10^-7^. Scale bars= 50 μm. **g**, Oxygen consumption rate measured during Rest, Dep, and DHPG stimulation in control and eIF4G2 knockdown neurons (n= 3 biological replicates). Dendritic mitochondrial mass and membrane potential (activity) measured in eIF4G2 WT, KD, and rescue conditions expressing dendritically targeted WT eIF4G2, dendritic T507V mutant, or somatic eIF4G2 in **h**, KCl-depolarized (Dep) neurons: Mass (WT vs. KD: p_adj_= 5.1x10^-4^; KD vs. KD+WT: p_adj_= 6.3x10^-3^; KD+WT vs. KD+T507V: p_adj_= 1.4x10^-3^, Potential (WT vs. KD: p_adj_= 1.3x10^-4^; KD vs. KD+WT: p_adj_= 1.5x10^-3^; KD+WT vs. KD+T507V: p_adj_= 5.1x10^-3^) and **i**, DHPG-activated neurons: Mass (WT vs. KD: p_adj_= 3.0x10^-2^, Potential (WT vs. KD: p_adj_= 1.4x10^-3^; KD vs. KD+WT: p_adj_= 1.1x10^-2^; KD+WT vs. KD+T507V: p_adj_= 4.0x10^-2^). **h**, **i**, Data are presented as mean ± SD. **f, h, i**, Statistical significance was determined by one-way ANOVA followed by Tukey’s multiple comparisons test.

To determine how dendritic PL-CLIP relates to conventional whole-cell eIF4G2 CLIP^12^, we compared normalized eIF4G2 binding across transcripts detected by both approaches under depolarizing conditions. Activity-dependent binding measured by PL-CLIP positively correlated with conventional CLIP (Spearman’s ρ = 0.45, p_val_= 5.7x10^-36^; Fig. 3c), indicating that dendritic PL-CLIP recapitulates a substantial fraction of canonical eIF4G2 targets while selectively resolving RNA interactions within postsynaptic compartments. These findings establish PL-CLIP as a compartment-resolved extension of conventional CLIP for mapping activity-dependent eIF4G2-RNA interactions specifically within dendrites.

We next examined the compartment-specific RNA targets identified by PL-CLIP (Supplementary Table 1). Activity-dependent eIF4G2 binding occurred predominantly within the 5′UTRs of transcripts whose coding sequences subsequently underwent increased dendritic translation (Fig. 3d,e and Extended Data Fig. 4c). Although eIF4G2 binding was also detected within other transcript regions, including 3′UTRs, activity-dependent enrichment was most pronounced at 5′UTRs, consistent with the established role of eIF4G2 in translation initiation (Extended Data Fig. 4c). Gene set enrichment analysis identified mitochondrial pathways among the most significantly enriched eIF4G2 targets (Fig. 3d and Extended Data Fig. 4d). Consistent with this result, the 5’UTRs of MitoCarta-annotated transcripts exhibited significantly greater activity-dependent eIF4G2 binding than non-mitochondrial transcripts following both KCl depolarization and DHPG stimulation (Extended Data Fig. 4e,f), demonstrating preferential recruitment of mitochondrial RNAs into the activated dendritic eIF4G2 interactome. Furthermore, transcripts identified by dendritic PL-CLIP underwent significantly greater activity-dependent CDS translation than either the transcriptome as a whole or transcripts identified by conventional whole-cell CLIP, as measured by PL-Ribo-seq (Fig. 3e and Extended Data Fig. 4g). Together, these findings indicate that eIF4G2 binding within dendrites selectively marks RNAs that subsequently undergo local translation during neuronal activation.

Among these targets, *Nsun3* emerged as a compelling candidate because it encodes a nuclear-encoded mitochondrial tRNA methyltransferase whose 5′UTR harbors an activity-responsive uORF and exhibits robust eIF4G2 binding following neuronal activation (Extended Data Fig. 5a). Notably, this interaction was not detected in our previous whole-cell eIF4G2 CLIP dataset^12^, suggesting that compartment-resolved PL-CLIP can identify dendritic interactions that are not readily detected at the whole-cell level. PCR amplification followed by Sanger sequencing independently validated selective recovery of the *Nsun3* 5′UTR from activated eIF4G2 PL-CLIP libraries (Extended Data Fig. 5b,c). Because *Nsun3* integrates multiple features of the eIF4G2 translational program, including activity-dependent 5′UTR binding, uORF translation, and mitochondrial protein synthesis, it provided an ideal model to investigate how eIF4G2 coordinates canonical and non-canonical translation during synaptic activity.

Following these observations, we asked whether activity-dependent eIF4G2 binding is required for the functional remodeling of dendritic mitochondria. We depleted eIF4G2 in primary neurons and quantified Δψm using the JC-1 ratiometric assay under resting and depolarized conditions. Control neurons exhibited the expected activity-dependent increase in Δψm, whereas this response was abolished following eIF4G2 knockdown (Fig. 3f), demonstrating that eIF4G2 is required for activity-induced enhancement of mitochondrial membrane potential.

Because mitochondrial membrane potential provides the proton-motive force required for oxidative phosphorylation, we next asked whether loss of eIF4G2 also affects mitochondrial respiration. Oxygen consumption increased following KCl depolarization and DHPG stimulation in control neurons but was markedly attenuated by either eIF4G2 knockdown or calcium chelation (Fig. 3g and Extended Data Fig. 6). These findings demonstrate that both calcium signaling and eIF4G2 are required to support the activity-dependent increase in mitochondrial respiratory capacity.

To determine whether eIF4G2 regulates mitochondrial responses to activity through a local dendritic mechanism, we performed compartment-specific rescue experiments using our dendritic targeting strategy (Fig. 2e). Following eIF4G2 knockdown, eIF4G2 was re-expressed either selectively in dendrites or as a somatically restricted construct (Fig. 3h and Extended Data Fig. 7). Dendritic eIF4G2 restored KCl-induced increases in mitochondrial mass and membrane potential, whereas the somatically restricted construct failed to rescue either phenotype (Fig. 3h). Following DHPG stimulation, dendritic eIF4G2 also restored membrane potential and partially rescued mitochondrial mass, although the latter did not reach statistical significance; neither effect was observed with somatic eIF4G2 (Fig. 3i). Furthermore, a dendritically localized phospho-deficient eIF4G2 T507V mutant, previously shown to disrupt activity-dependent translational induction^12^, was also ineffective. Together, these findings support that activity-responsive eIF4G2 is locally required for dendritic mitochondrial adaptation and that its dendritic re-expression is sufficient, with the magnitude of the effect varying across stimuli and mitochondrial readouts.

### eIF4G2 coordinates canonical protein synthesis and uORF translation in dendrites

We previously showed that eIF4G2 preferentially associates with the 5′UTRs of activity-responsive transcripts to promote downstream translation^12^. Many of these 5′UTRs contain uORFs, which are increasingly recognized as dynamic regulators of translation and, in some cases, as sources of functional micropeptides^35,44–47^. To determine whether activity-dependent dendritic eIF4G2 binding preferentially targets this class of transcripts, we compared ORF-RATER-derived uORF scores between eIF4G2-bound and unbound dendritic RNAs. eIF4G2-bound transcripts exhibited significantly higher uORF scores (Fig. 4a), suggesting that eIF4G2 preferentially recognizes transcripts capable of producing both canonical proteins and uORF-encoded peptides during neuronal activation. Representative PL-Ribo-seq profiles further showed activity-dependent ribosome occupancy across uORFs in several dendritic transcripts, including *Nsun3* (Fig. 4b).

**Fig. 4.**
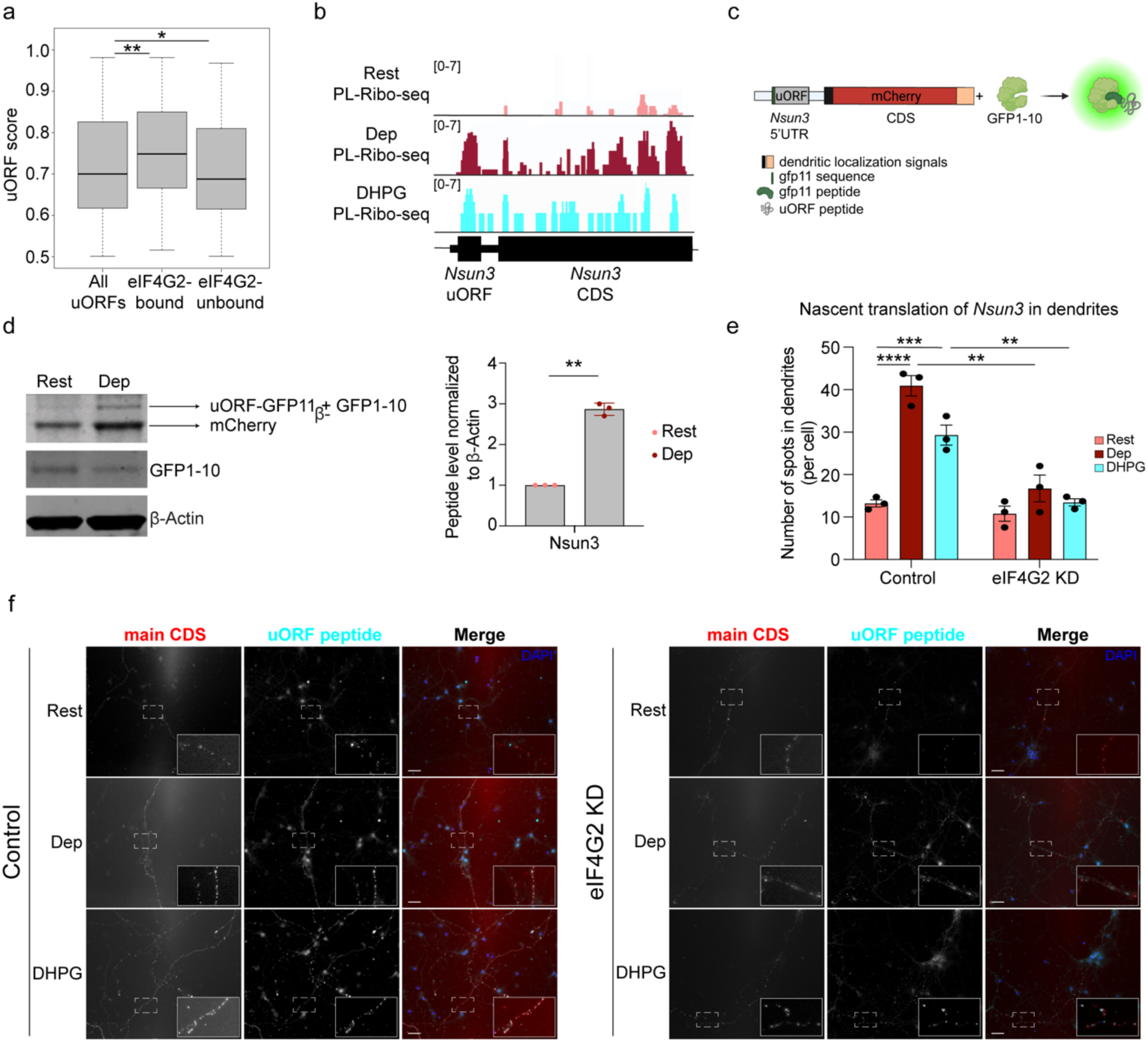
uORFs in dendritic transcripts encode micropeptides that are upregulated by synaptic activity and eIF4G2. a, ORF-RATER-derived uORF scores, grouped according to eIF4G2 binding status of the uORF-containing 5′UTR. Statistical significance was assessed using a two-sided paired Wilcoxon signed-rank test: All vs. eIF4G2-bound: p_adj_= 7.8x10^-3^ ; All vs. eIF4G2-unbound: p_adj_= 1.3x10^-2^). **b**, Representative PL-Ribo-seq tracks showing ribosome occupancy over uORFs in dendritic transcripts, including *Nsun3*, under resting conditions and following neuronal activation by depolarization or DHPG treatment. **c**, Schematic of the dual-reporter assay used to monitor uORF-encoded micropeptide translation in neurons. The uORF sequence is fused in-frame to GFP11 and co-expressed with GFP1-10, enabling fluorescence complementation upon micropeptide translation. **d**, Native gel immunoblots detecting complemented GFP signal from the *Nsun3* uORF reporter, together with GFP1-10 expression and β-actin as a loading control. Quantification shows complemented GFP signal normalized to β-actin under resting and depolarized conditions. Statistical significance was determined by two-tailed paired Student’s *t*-test. **e**, Quantification of puro-PLA signal detecting nascent *Nsun3* translation in dendrites of control siRNA-and eIF4G2 siRNA-treated neurons under resting and activated conditions (n= 3 biological replicates). Control Rest vs. Dep: padj= 1.9x10^-6^; Control Rest vs. DHPG: p_adj_= 3.8x10^-4^; Control Dep vs. KD Dep: p_adj_= 1.8x10^-3^; Control DHPG vs. KD DHPG: p_adj_= 1.6x10^-3^. Data are presented as mean ± SD. Statistical significance was determined by one-way ANOVA followed by Tukey’s multiple comparisons test. **f**, Representative images and quantification of uORF-encoded micropeptide and main CDS protein signals from the reporter shown in **(c)** under resting and activated conditions after control or eIF4G2 siRNA treatment. Scale bars= 50 μm.

To test directly whether these uORFs could produce detectable peptides in neurons, we developed a split-GFP complementation reporter in which individual uORFs were fused in frame to GFP11 and co-expressed with GFP1-10 (Fig. 4c). GFP fluorescence therefore depended on translation of the GFP11-tagged uORF product. Fusion of GFP11 to the β-actin coding sequence produced robust fluorescence complementation, confirming that the reporter system could detect translated GFP11-tagged peptides (Extended Data Fig. 8).

We next applied this strategy to the activity-responsive *Nsun3* uORF. Neuronal activation increased GFP complementation from the *Nsun3* uORF reporter, demonstrating that the uORF is translated into a detectable peptide following stimulation (Fig. 4d,f). These findings provide direct evidence that activity-dependent dendritic translation generates previously unannotated micropeptides from neuronal uORFs.

We then asked whether activity-dependent translation of the *Nsun3* uORF and downstream coding sequence depended on eIF4G2. eIF4G2 knockdown reduced translational upregulation across both the endogenous *Nsun3* 5′UTR and coding sequence, as measured by dendritic PL-Ribo-seq (Extended Data Fig. 9a). Consistent with this result, activity-induced nascent NSUN3 synthesis measured by puro-PLA was strongly reduced following eIF4G2 depletion under both KCl and DHPG stimulation (Fig. 4e and Extended Data Fig. 9b,c). In parallel, eIF4G2 knockdown diminished both uORF-derived GFP complementation and downstream reporter expression (Fig. 4f). By contrast, activity-dependent translation of Tspo, which was not detectably bound by eIF4G2, remained unchanged following eIF4G2 depletion, providing an internal control for the specificity of this regulatory pathway (Extended Data Fig. 9d).

Together, these findings demonstrate that eIF4G2 is required for activity-dependent translation of both the Nsun3 uORF and downstream coding sequence in dendrites, while leaving a non-target mitochondrial transcript unaffected. Thus, activity-dependent dendritic eIF4G2 coordinates the local synthesis of both mitochondrial proteins and previously unannotated uORF-encoded micropeptides, revealing an unexpected layer of synaptic gene regulation.

### Local translation of NSUN3 mediates activity-dependent mitochondrial remodeling

Having established *Nsun3* as a dendritic eIF4G2 target and shown that its activity-dependent translation requires eIF4G2, we next asked whether locally synthesized NSUN3 affects mitochondrial function. To test this, we depleted *Nsun3* and measured dendritic mitochondrial mass and membrane potential under resting and activation conditions. Both parameters increased after activation in control neurons, whereas these responses were impaired following *Nsun3* depletion (Fig. 5a,b). The effects under resting conditions were more modest and not significant, suggesting that NSUN3 is required for the mitochondrial adaptation that accompanies neuronal activity rather than for maintenance of basal mitochondrial properties.

**Fig. 5.**
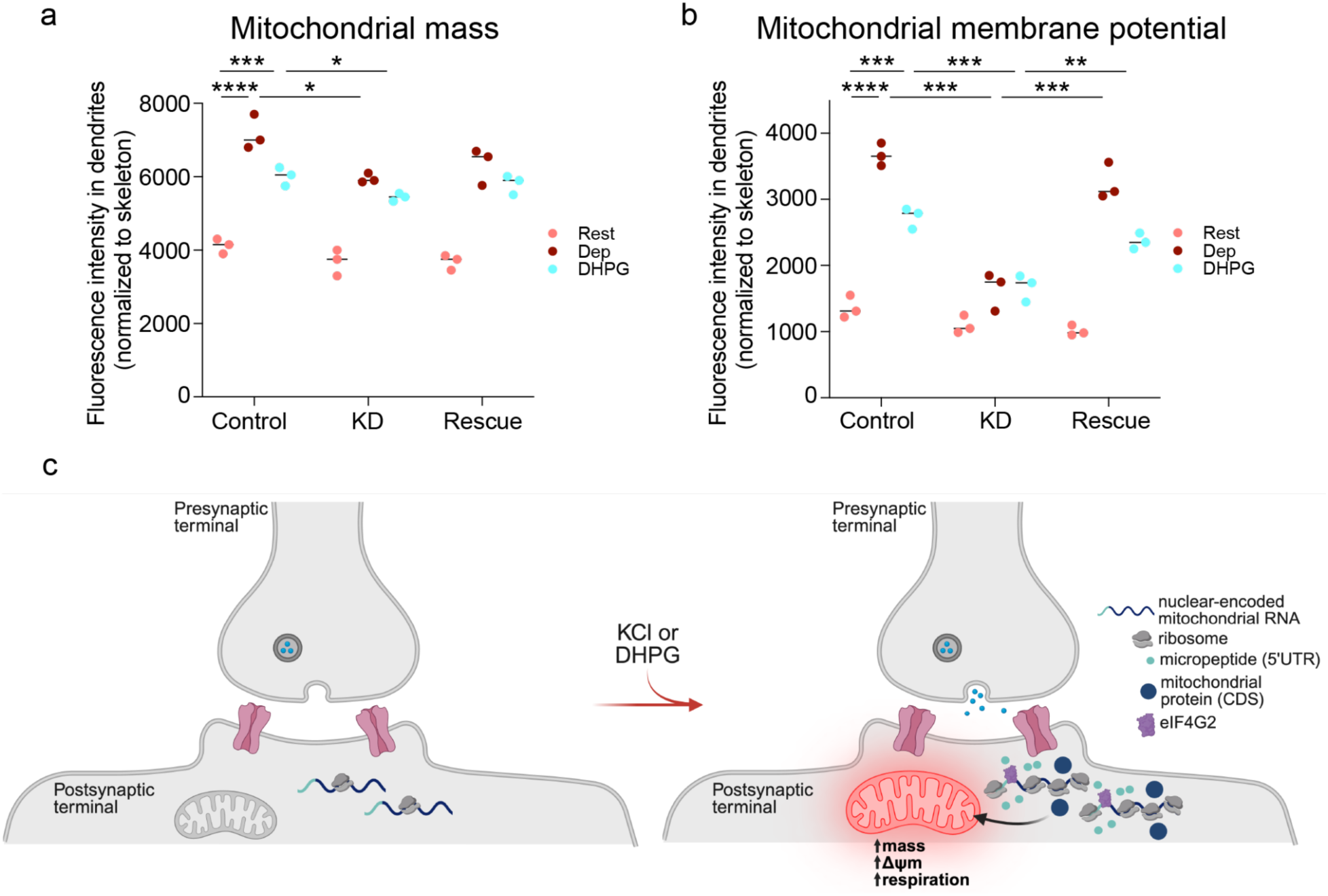
eIF4G2-directed translation of synaptic mitochondrial RNAs remodels local dendritic mitochondria. Quantification of **a**, mitochondrial mass and **b**, mitochondrial membrane potential, measured by MitoTracker Green FM and MitoTracker Deep Red FM fluorescence, respectively, in dendrites of control, NSUN3 knockdown (KD), and dendritic NSUN3 rescue neurons under resting conditions or following KCl depolarization (Dep) or DHPG stimulation. Dendritic rescue was achieved using the localization strategy shown in Fig. 2e. Fluorescence intensity was normalized to dendritic skeleton length. Median values are shown (n= 3 biological replicates). Statistical significance was determined by one-way ANOVA followed by Tukey’s multiple comparisons test. Mitochondrial mass: Rest vs. Dep Control p_adj_= 7.1x10^-5^; Rest vs. DHPG Control p_adj_= 9.9x10^-4^; Dep Control vs. Dep KD p_adj_= 2.4x10^-2^; DHPG Control vs. DHPG KD p_adj_= 4.2x10^-2^. Mitochondrial activity: Rest vs. Dep Control p_adj_= 6.7x10^-6^; Rest vs. DHPG Control p_adj_= 1.4x10^-4^; Dep Control vs. Dep KD p_adj_= 1.0x10^-4^; DHPG Control vs. DHPG KD p_adj_= 5.0x10^-4^; Dep KD vs. Dep Rescue p_adj_= 5.5x10^-4^; DHPG KD vs. DHPG Rescue p_adj_= 5.2x10^-3^. **c,** In resting dendrites, nuclear-encoded mitochondrial RNAs are enriched in postsynapses but undergo limited local translation. Upon synaptic activation by KCl depolarization or DHPG stimulation, eIF4G2 is recruited to synaptic mRNAs and promotes local translation near dendritic mitochondria. This activity-induced translation produces mitochondrial regulatory proteins and uORF-encoded micropeptides, which together support remodeling of local mitochondria, including increased mitochondrial mass, membrane potential (activity), and function.

To determine whether local synthesis of NSUN3 is sufficient to restore mitochondrial response, we re-expressed dendritically targeted NSUN3 in knockdown neurons using the localization strategy described in Fig. 2e. Dendritic NSUN3 recovered activity-dependent mitochondrial membrane potential after KCl and DHPG stimulation and partially rescued mitochondrial mass, although this effect did not reach statistical significance (Fig. 5a,b). These findings demonstrate that local NSUN3 synthesis is sufficient to restore activity-dependent mitochondrial membrane potential, whereas its effect on mitochondrial mass remains less clear.

Together, these results support a model in which activity-evoked calcium signaling induces a local translational program in dendrites that is mediated, in part, by eIF4G2 and includes nuclear-encoded mitochondrial proteins and uORF-derived micropeptides. These locally translated proteins, in turn, modulate mitochondrial function and establish mitochondrial competence needed to sustain activity-dependent calcium signaling at synapses (Fig. 5c).

## Discussion

Synaptic activity imposes rapid and highly localized energetic demands that require localized adaptive responses in dendrites. Here, we identify a dendrite-specific translational mechanism that couples neuronal activity to mitochondrial regulation through the non-canonical initiation factor eIF4G2. By integrating compartment-resolved RNA interactome mapping with local translation profiling and functional analyses, we show that neuronal activation increases eIF4G2 binding to a subset of nuclear-encoded mitochondrial transcripts and promotes their translation locally. This response was accompanied by increases in mitochondrial membrane potential, abundance, and respiration. These findings demonstrate that local translation is not simply a mechanism to alter available proteins at a synapse but can also adapt the state of nearby mitochondria to neuronal activity.

An important finding of this study is that there is reciprocal regulation between calcium signaling, local translation, and mitochondrial function in dendrites. Previous studies showed that synaptic calcium entry controls mitochondrial positioning through the calcium-sensitive trafficking factor Miro1 and that PDZD8-mediated contacts between ER and mitochondria enable mitochondrial calcium uptake and shape dendritic calcium dynamics^41,42^. We identify localized translation as a molecular mechanism that couples these two directions. Calcium influx stimulates local translation, and some of the locally synthesized mitochondrial proteins reinforce the mitochondrial competence needed to sustain dendritic calcium signaling. Local protein synthesis may therefore help mitochondria respond to the energetic demands of synaptic activation and maintain subsequent persistent signaling. In line with this idea, depletion of TSPO or NSUN3 impaired calcium responses triggered by both depolarization and mGluR activation, and dendritic re-expression restored them. Recent work showing that synaptic calcium signals acutely stimulate local mitochondrial ATP synthesis to support plasticity is consistent with a reciprocal relationship between mitochondrial energetic state and synaptic signaling^48^. Our data therefore suggest that changes in mitochondrial function are not only a response to calcium influx but also help maintain calcium signaling after neuronal activation. Rather than operating as a linear pathway, calcium signaling, local translation, and mitochondrial remodeling may form a positive feedback circuit that enables dendrites to maintain both bioenergetic capacity and signaling competence during sustained synaptic activation.

Our findings also expand the functional repertoire of eIF4G2. Previous studies established eIF4G2 as a non-canonical translation initiation factor capable of regulating transcripts containing structured 5′UTRs and uORFs^49,50^. By resolving dendrite-specific eIF4G2-RNA interactions, we found that its activity-dependent binding is both spatially and temporally regulated, preferentially targeting transcripts that subsequently undergo local translation. Opto-CLIP, a related approach combining optogenetic stimulation with cell-type-specific CLIP, similarly enables activity-dependent RBP-RNA interactions to be resolved in defined neuronal populations^51^. However, what modulates eIF4G2 upstream still needs to be investigated. Activity-dependent eIF4G2 enrichment was most prominent within 5′UTRs, consistent with its established role in translation initiation^49,52^. However, PL-CLIP also detected eIF4G2 binding within 3′UTRs, raising the possibility that eIF4G2 participates in additional regulatory processes beyond translation initiation. Such interactions could contribute to mRNP remodeling, transcript circularization, or coordination of translation with 3′UTR-dependent mechanisms governing RNA localization, stability, or translational efficiency. In particular, mGluR activation elicited a distinct dendritic translational program characterized by increased eIF4G2 association with RNAs involved in membrane signaling, calcium regulation, intracellular transport, and mitochondrial and metabolic adaptation. This selective binding was accompanied by preferential translational activation of mitochondrial transcripts and other RNAs associated with 3′UTR-dependent regulation, suggesting that eIF4G2 orchestrates specialized local translational responses rather than globally enhancing protein synthesis. One intriguing possibility is that local synthesis of membrane signaling components, including G-protein signaling effectors identified among DHPG-responsive transcripts, reinforces or prolongs mGluR signaling through a positive feedback mechanism.

Our data further suggest that eIF4G2 is not simply permissive for translation but instead licenses an activity-dependent translational switch in dendrites. Transcripts that showed increased eIF4G2 binding after neuronal activation were also more likely to undergo local translation. Accordingly, eIF4G2 depletion prevented activity-dependent translation of both the *Nsun3* uORF and its downstream coding sequence and altered the increase in uORF ribosome occupancy normally observed after stimulation. One possibility is that eIF4G2 stabilizes initiation complexes on structured 5′UTRs or promotes productive ribosome engagement at both uORFs and downstream coding sequences. It may also facilitate reinitiation after uORF translation, protect its target transcripts from translational repression, or help assemble specialized initiation complexes at active synapses. Previous work showed that phosphorylation of specific eIF4G2 residues is required for its activity-dependent functions^12^, but how these modifications influence initiation on individual target transcripts and which of these mechanisms underlies this regulation remain to be determined.

Our work also provides direct evidence that activity-dependent dendritic translation generates previously unannotated micropeptides from endogenous neuronal uORFs. Although uORFs have long been appreciated as regulators of downstream translation, their contribution to the local neuronal proteome has remained largely unexplored. By adapting split-GFP complementation to visualize translation of endogenous neuronal uORFs, we demonstrate that activity-dependent local translation expands the dendritic proteome beyond annotated coding sequences. Together with our observation that eIF4G2 coordinates translation of both uORFs and their downstream coding sequences, these findings reveal that eIF4G2 directs a dual-output translational program in which a single dendritic transcript gives rise to both a mitochondrial protein and an activity-regulated micropeptide. More broadly, this work uncovers a vast and previously unexplored layer of synaptic gene regulation and raises the possibility that numerous activity-regulated uORFs identified in dendrites encode functional micropeptides that participate in synaptic signaling, mitochondrial biology, and neuronal plasticity. This work therefore opens a path toward defining the functions of this previously uncharacterized micropeptide proteome and determining how its activity-dependent production shapes neuronal signaling, organelle function, and synaptic plasticity.

Together, our findings establish localized translation as a mechanism for coordinating organelle adaptation with neuronal activity. Rather than functioning solely to replenish synaptic proteins, dendritic translation actively remodels mitochondrial physiology, allowing individual synapses to rapidly adapt their metabolic capacity to local signaling demands. Given the widespread disruption of both mitochondrial function and local translation across neurodevelopmental and neurodegenerative disorders, this activity-dependent regulatory framework may represent a fundamental mechanism through which neurons maintain synaptic function and resilience.

## Acknowledgements

This work was supported by NIH grant R35NS097404 (NINDS) awarded to R.B.D. and NIH grant 5R35GM159898 (NIGMS) awarded to E.H. R.B.D. is an Investigator of the Howard Hughes Medical Institute. E.H. also acknowledges startup support from the Wertheim UF Scripps Institute. We thank Robert M. Witwicki, Li Pan, and Marlene L. Biller of the Wertheim UF Scripps Institute Genomics Core for sequencing support and technical assistance. We are grateful to the members of the Hacisuleyman and Darnell laboratories for insightful discussions throughout this work. We especially thank R.A. Singer for valuable discussions and critical reading of the manuscript.

## Author contributions

R.B.D. and E.H. conceived and designed the study. M.T.J and E.H. administered the project and conducted all the experiments. M.T.J and E.H. performed all the analyses. N.N. contributed to imaging and reporter experiments. R.B.D. and E.H. obtained funding. M.T.J., R.B.D., and E.H. wrote and reviewed the manuscript. All authors have edited the manuscript.

## Data and materials availability

All sequencing data generated in this study have been deposited in Gene Expression Omnibus with the accession number GSE339407. All other data and code used in this study are available or described in the manuscript or supplementary materials.

## Extended Data Figures

**Extended Data Fig. 1.**
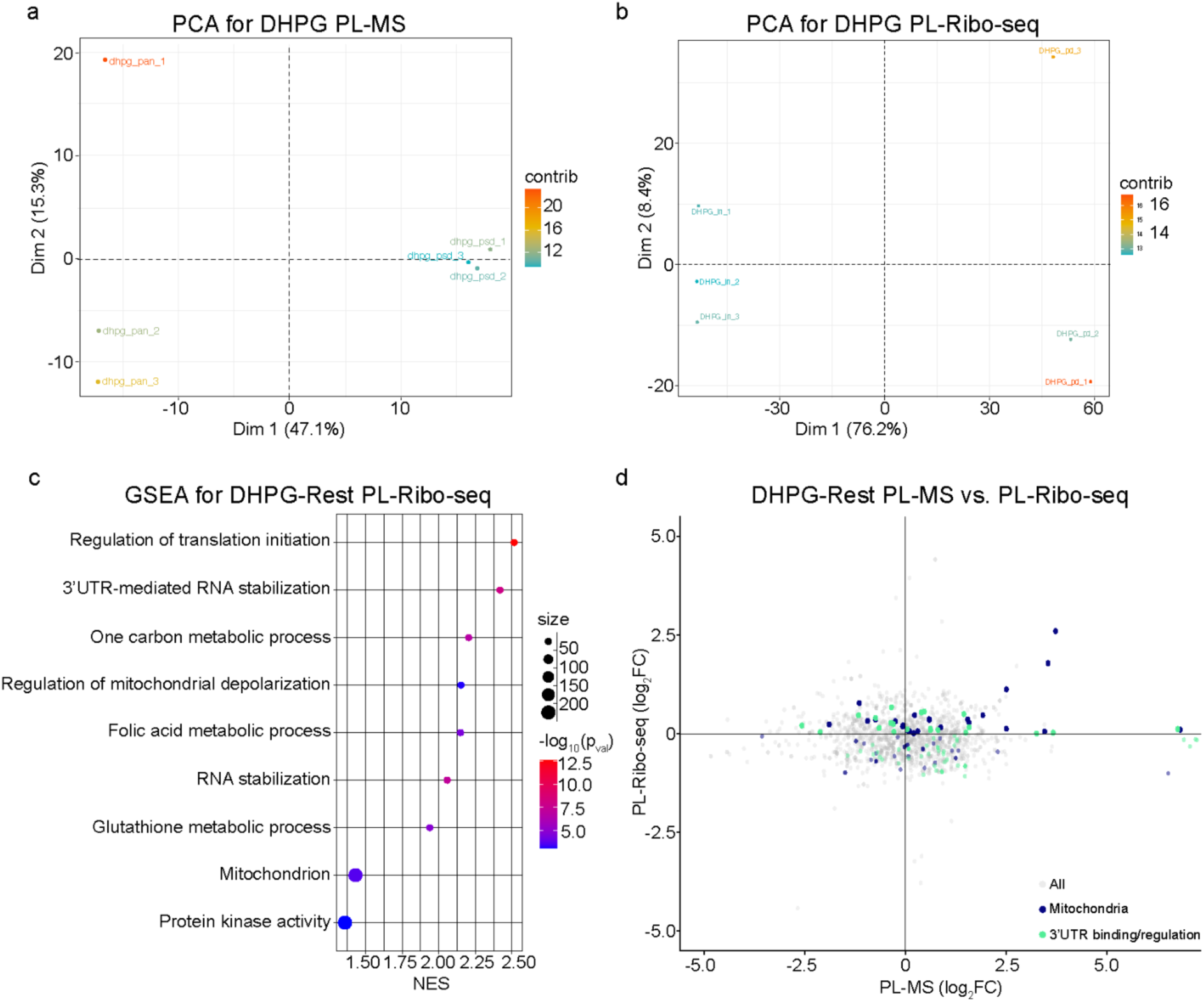
DHPG remodels dendritic proteomic and translational programs. **a**, PCA plot of three PL-MS biological replicates from DHPG-treated neurons transduced with Pan-TurboID or TurboID-PSD95. For each sample, the corresponding minus-biotin control was subtracted prior to plotting. **b**, PCA plot of three PL-Ribo-seq biological replicates from input and pulldown fractions of TurboID-PSD95-transduced neurons following DHPG stimulation. **c**, Gene set enrichment analysis (GSEA) of RNAs with increased dendritic translation upon DHPG treatment, as measured by PL-Ribo-seq differential analysis comparing DHPG versus resting conditions. Significantly enriched gene sets are shown at FDR < 0.05, with multiple-testing correction performed using the Benjamini-Hochberg method. **d**, Distribution of mitochondria-related and 3′ UTR regulation-related RNAs across differential PL-MS and PL-Ribo-seq datasets comparing DHPG-treated and resting neurons.

**Extended Data Fig. 2.**
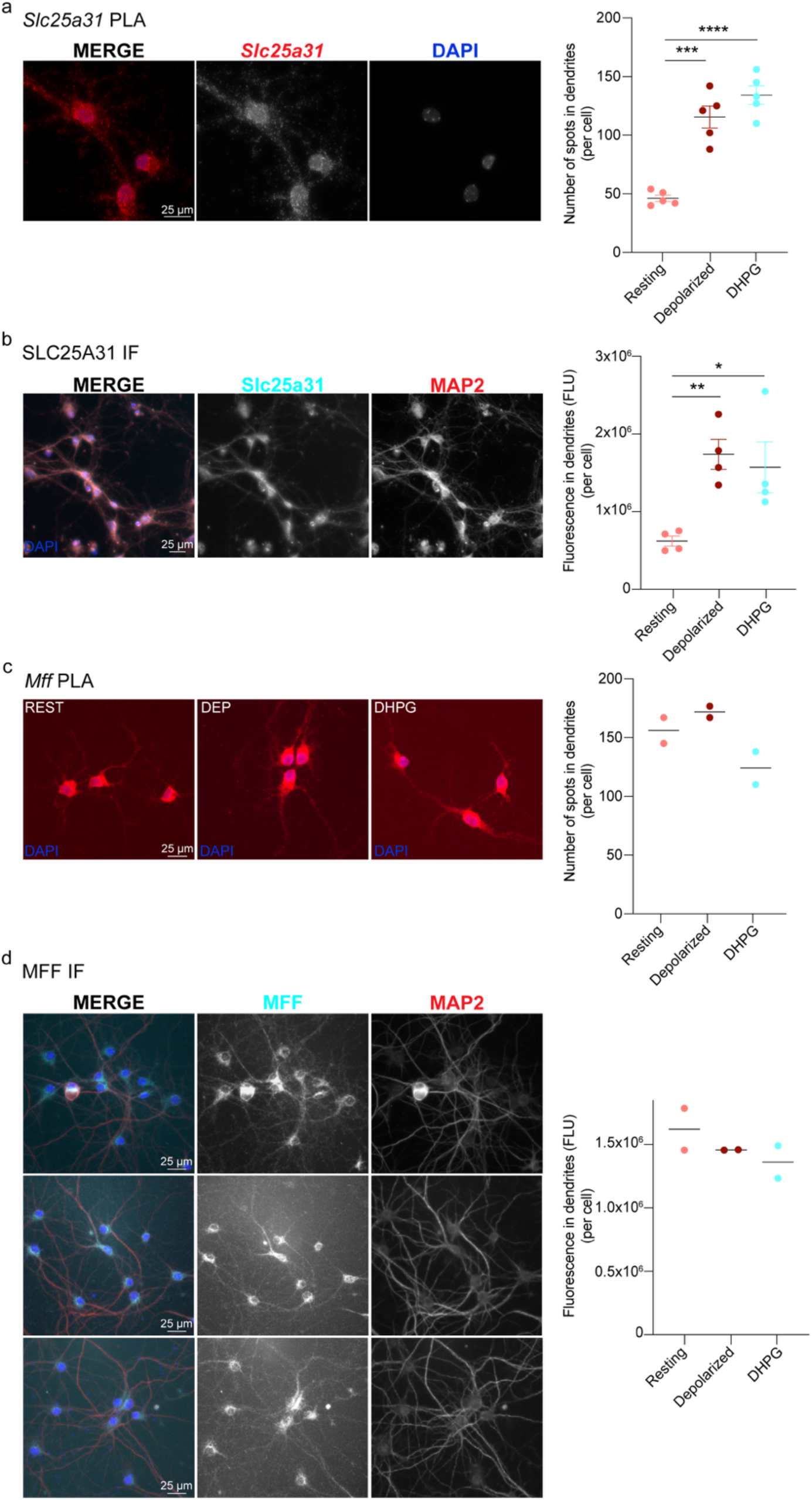
DHPG stimulation selectively increases dendritic Slc25a31 translation and SLC25A31 protein abundance. In addition to the resting and KCl-depolarized conditions presented in Fig. 1d,e, DHPG-treated counterparts are shown for **a**, nascent *Slc25a31* translation measured by puromycin proximity ligation assay (puro-PLA; n= 5 biological replicates; Rest vs. DHPG, p_adj_= 5.4x10^-6^) and **b**, total SLC25A31 protein levels measured by immunofluorescence (IF; n= 4 biological replicates; Rest vs. DHPG, p_adj_= 3.0x10^-2^), together with their corresponding quantifications. Three fields were imaged per biological replicate. Data are presented as mean ± SEM. Statistical significance was determined by one-way ANOVA followed by Tukey’s multiple-comparisons test. **c,** Representative images and quantification of nascent dendritic *Mff* translation measured by puro-PLA under resting, KCl-depolarized and DHPG-treated conditions (n= 2 biological replicates). **d,** Representative images and quantification of dendritic MFF protein abundance measured by IF under the same conditions (n= 2 biological replicates). Three fields were imaged per biological replicate. In **a,b**, data are presented as mean ± SEM, and statistical significance was determined by one-way ANOVA followed by Tukey’s multiple-comparisons test. In **c,d**, horizontal lines indicate the mean. Scale bars= 25 μm.

**Extended Data Fig. 3.**
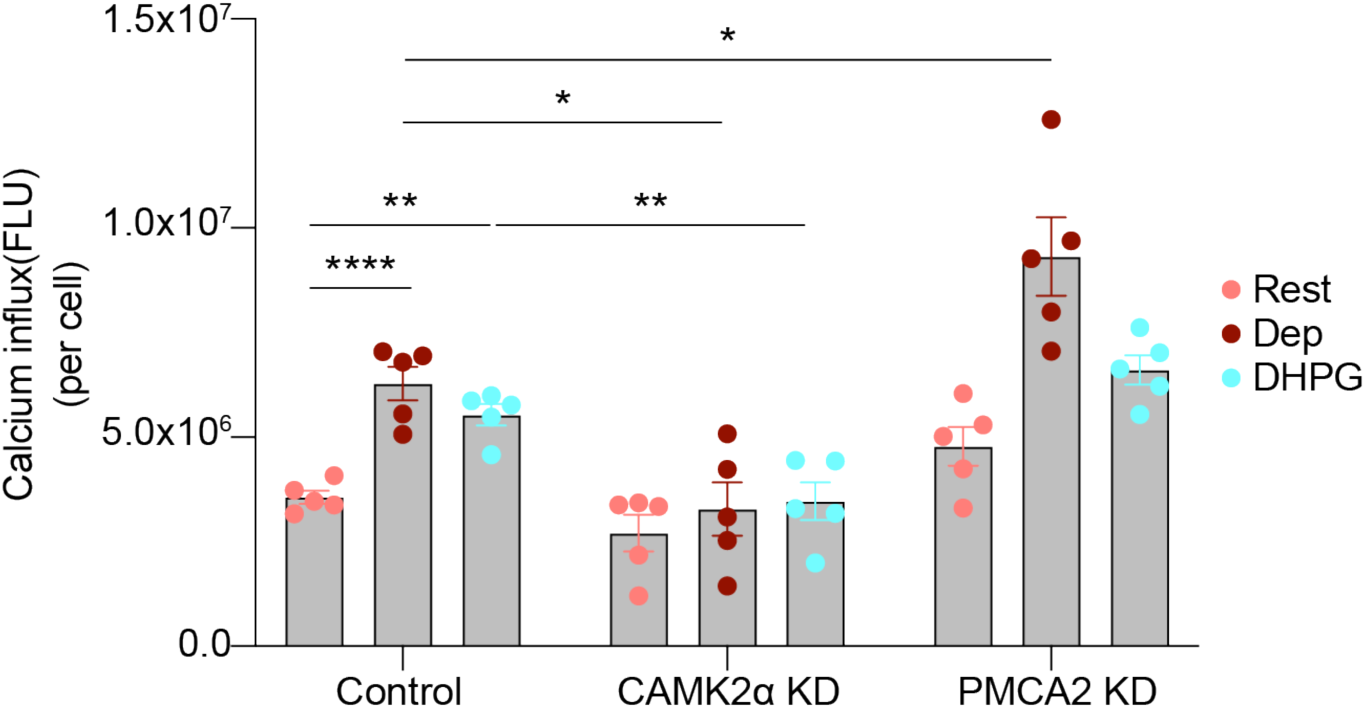
CAMK2A and PMCA2 bidirectionally regulate neuronal calcium responses. Quantification of Fluo-4 AM fluorescence in control, CAMK2α knockdown (KD), and PMCA2 KD neurons under resting (Rest), KCl depolarization (Dep), and DHPG stimulation conditions. CAMK2A depletion reduced, whereas PMCA2 depletion increased, stimulus-evoked intracellular calcium levels. Data are presented as mean ± SEM with individual biological replicates shown, and statistical significance was determined by one-way ANOVA followed by Tukey’s multiple-comparisons test (n= 5 biological replicates). Control Rest vs. Dep: p_adj_= 6.8x10^-5^; Control Rest vs. DHPG: p_adj_= 1.2x10^-3^; Control Dep vs. CAMK2α KD Dep: p_adj_= 2.5x10^-2^; Control DHPG vs. CAMK2α KD DHPG: p_adj_= 4.5x10^-3^; Control Dep vs. PMCA2 KD Dep: p_adj_= 2.4x10^-2^.

**Extended Data Fig. 4.**
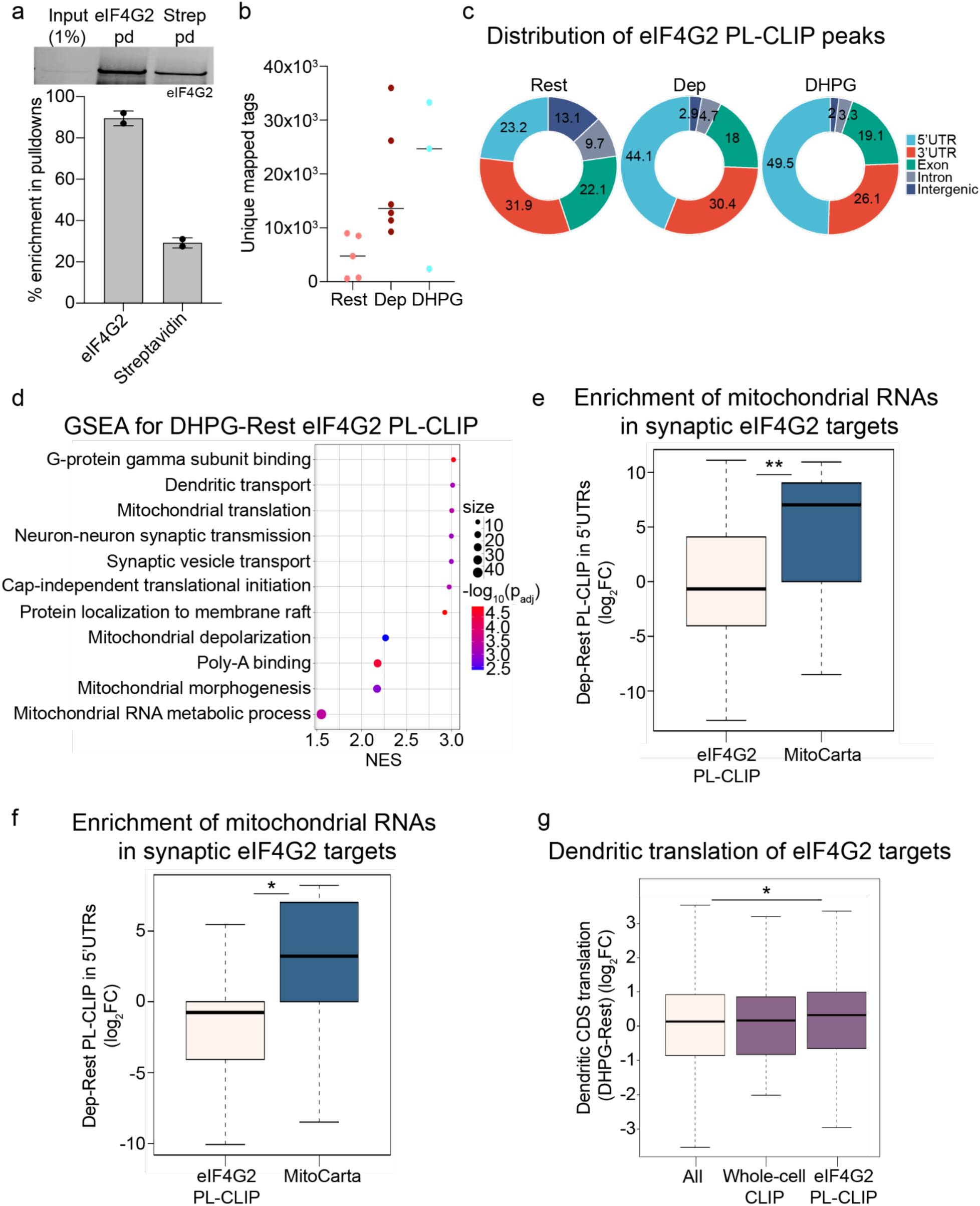
PL-CLIP identifies activity-regulated dendritic targets of eIF4G2. **a**, Western blot and quantification showing eIF4G2 enrichment after sequential eIF4G2 immunoprecipitation and streptavidin pulldown in the PL-CLIP protocol. Data are presented as mean ± SD. **b**, Number of eIF4G2 PL-CLIP tags uniquely mapped to the mouse genome (mm10) following removal of duplicates in resting, KCl-depolarized (Dep), and DHPG-stimulated libraries. **c,** Distribution of eIF4G2 PL-CLIP peaks across annotated transcript regions in resting, KCl-depolarized and DHPG-treated neurons, shown as the percentage of peaks mapping to each transcript feature. **d**, Gene set enrichment analysis (GSEA) of RNAs differentially bound by eIF4G2 in DHPG-treated compared with resting neurons, as identified by PL-CLIP. Significantly enriched gene sets are shown at FDR < 0.05, with multiple-testing correction performed using the Benjamini-Hochberg method. **e,f,** Distribution of activity-dependent eIF4G2 PL-CLIP binding changes for MitoCarta-annotated transcripts following **e,** KCl depolarization (Dep-Rest; p_val_= 1.5x10^-3^) and **f,** DHPG stimulation (DHPG-Rest; p_val_= 3.3x10^-2^). **g**, Dendritic CDS translation measured by PL-Ribo-seq, shown as DHPG-Rest log₂ fold change, for all RNAs and for eIF4G2-bound RNAs identified by whole-cell CLIP or eIF4G2 PL-CLIP (p_val_= 1.5x10^-2^). **e-g**, Box plots show the median, interquartile range and whiskers extending to 1.5x the interquartile range. Statistical significance was assessed using a two-sided paired Wilcoxon signed-rank test.

**Extended Data Fig. 5.**
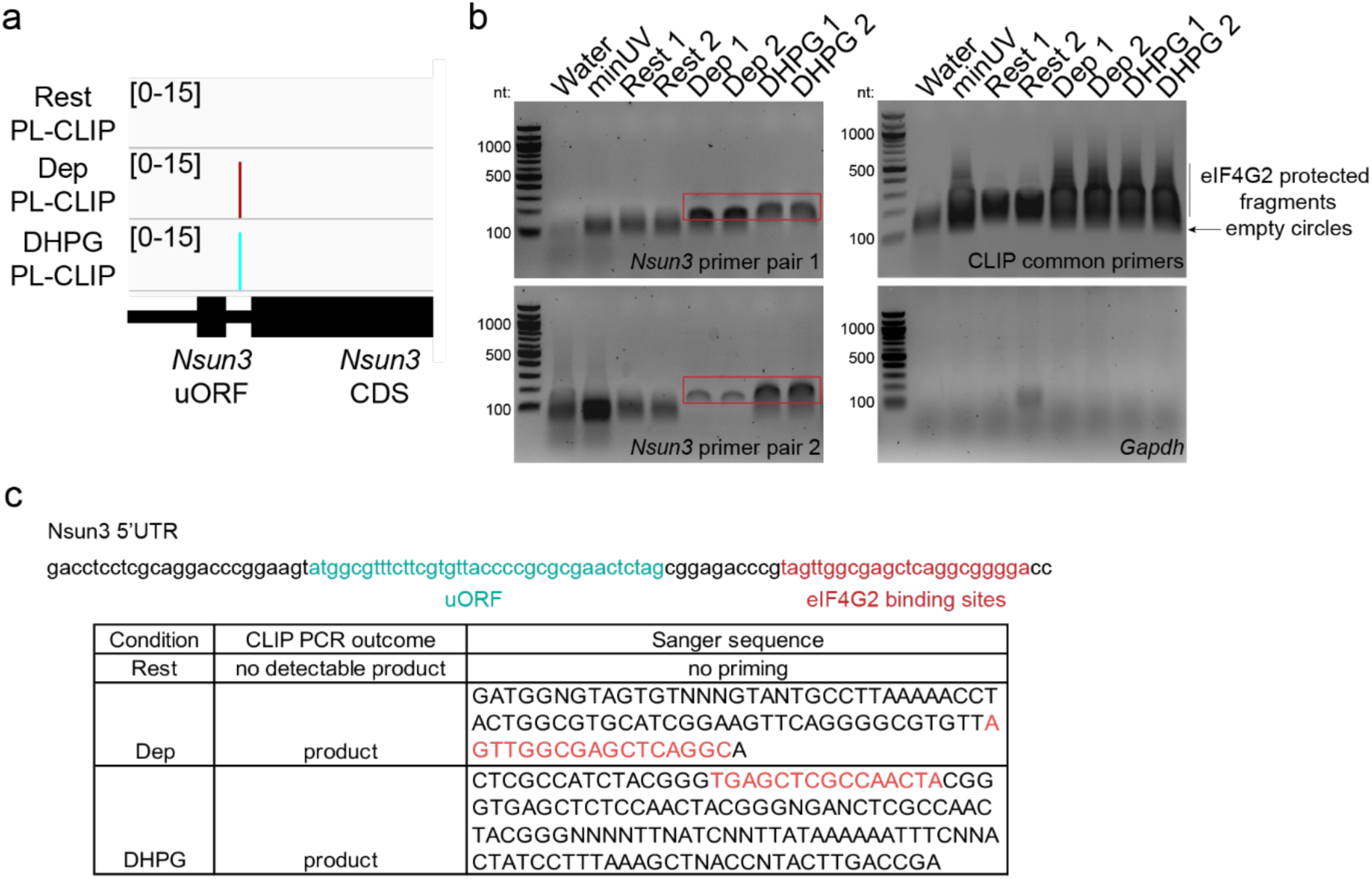
eIF4G2 PL-CLIP detects activity-dependent binding to the *Nsun3* 5′ UTR. a, Genome browser tracks (peak heights, log_2_) showing eIF4G2 proximity-labeled CLIP signal across the *Nsun3* 5′ UTR in resting, KCl-depolarized, and DHPG-treated neurons. The *Nsun3* uORF and eIF4G2-bound region are indicated. **b,** PCR validation of *Nsun3* 5′ UTR recovery from eIF4G2 PL-CLIP libraries using two independent *Nsun3* primer pairs. Water and minus UV samples serve as negative controls. CLIP common primers targeting linker and barcode-containing products confirm amplification of CLIP-derived fragments, whereas *Gapdh* serves as a negative control. PCRs were amplified using the same cycle number across conditions. **c,** Sanger sequencing of PCR products from activated conditions confirms that amplified fragments map to the *Nsun3* 5′ UTR, including the region containing the eIF4G2 binding sites. No detectable *Nsun3* PCR product was obtained from resting samples under the same amplification conditions.

**Extended Data Fig. 6.**
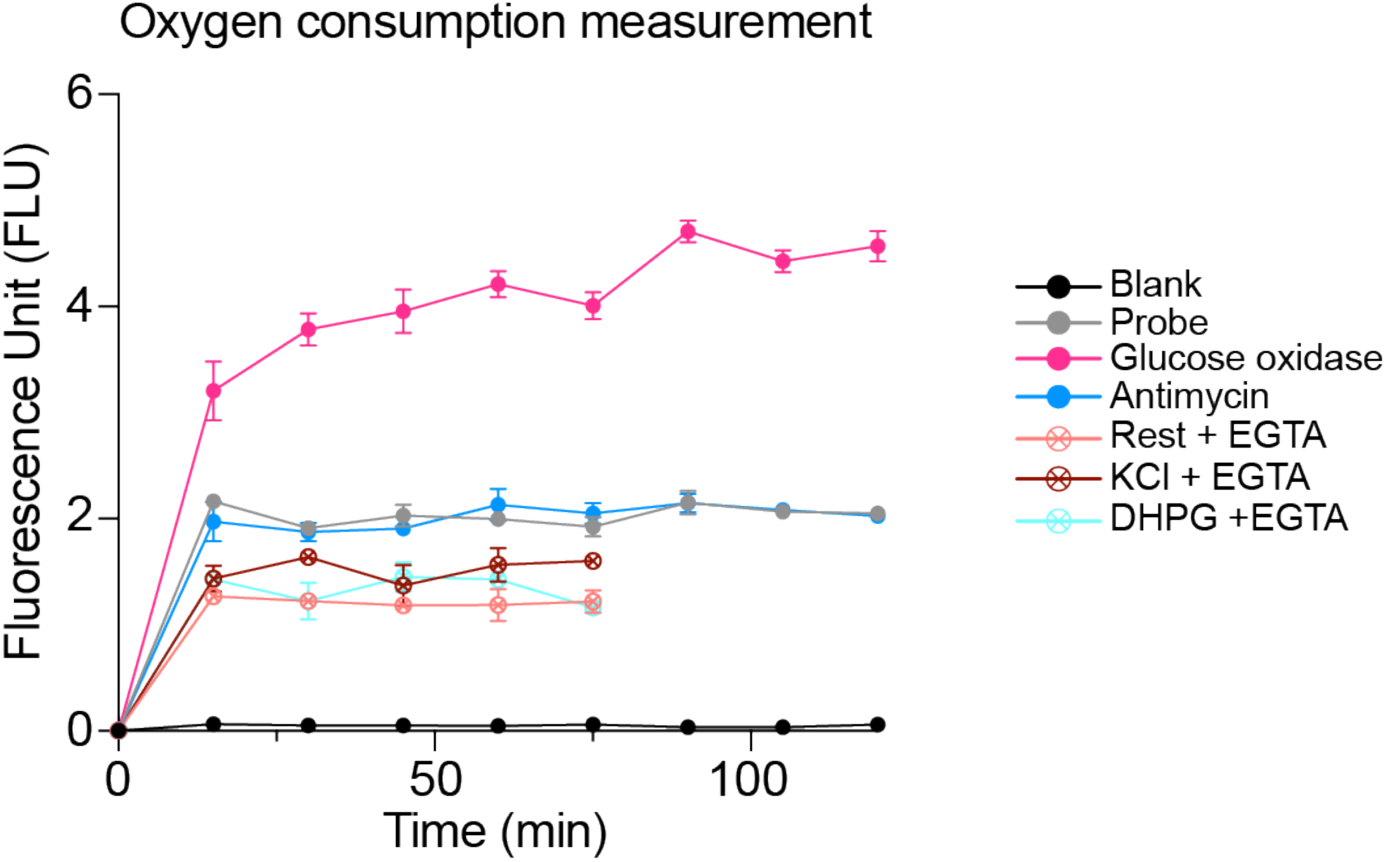
Assay controls validate calcium-dependent oxygen consumption measurements. Time-course of oxygen-sensitive probe fluorescence in blank wells, probe-only wells, glucose oxidase positive-control wells, antimycin-treated neurons, and neurons under resting, KCl-depolarized, or DHPG-treated conditions in the presence of EGTA. Glucose oxidase produced a robust increase in fluorescence, whereas antimycin-treated samples remained comparable to probe-only controls, confirming assay responsiveness and mitochondrial dependence. EGTA reduced the signal under resting, KCl-depolarized, and DHPG-treated conditions, supporting calcium-dependent regulation of activity-induced oxygen consumption. Data are presented as mean ± SD.

**Extended Data Fig. 7.**
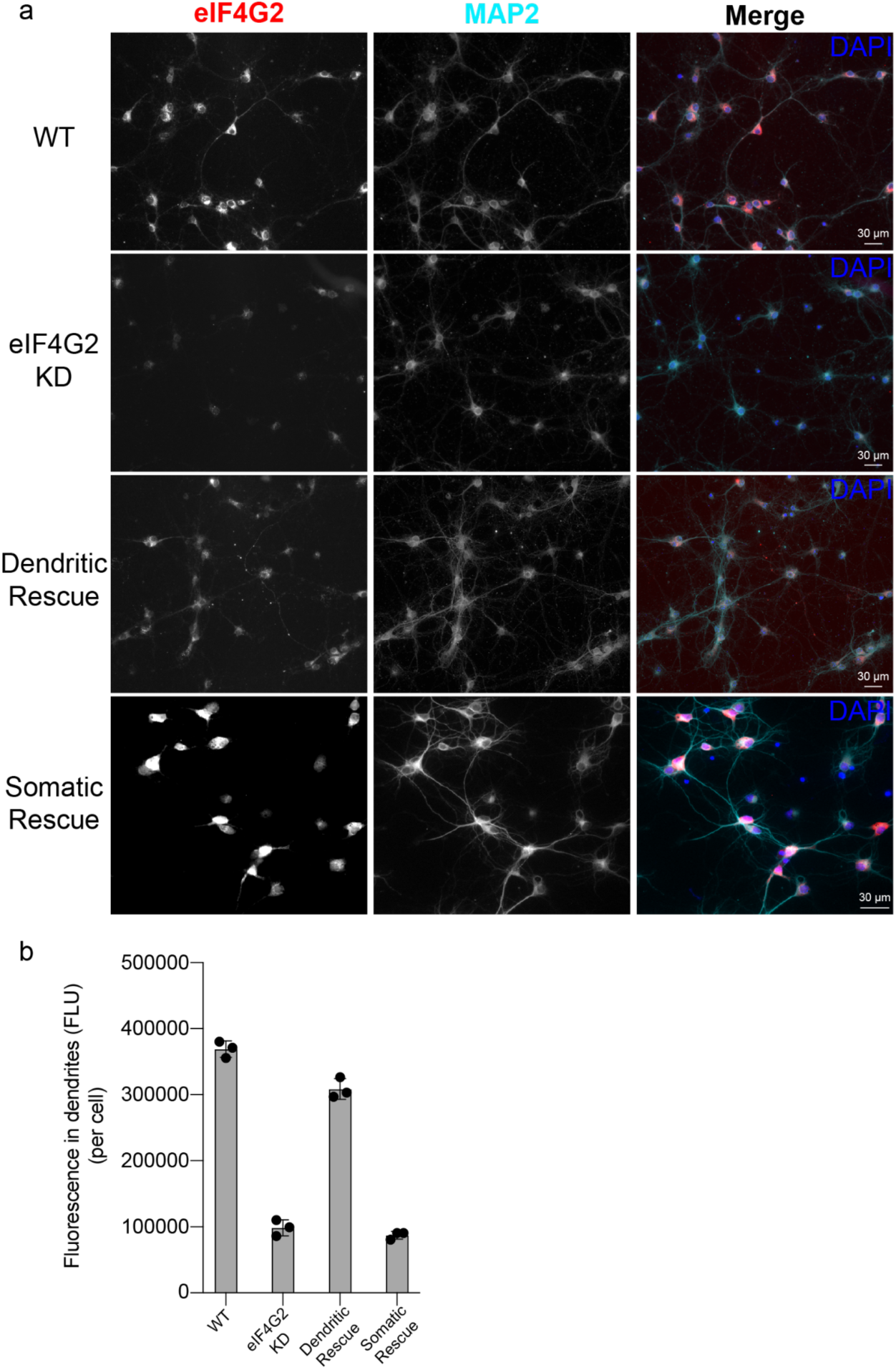
eIF4G2 immunofluorescence confirms knockdown and compartment-specific rescue in neurons. a, Representative IF images of primary cortical neurons stained for eIF4G2 (red), MAP2 (cyan), and DAPI (blue) under wild-type (WT), eIF4G2 knockdown (KD), dendritic rescue, and somatic rescue conditions. eIF4G2 immunoreactivity is markedly reduced following knockdown and restored following expression of the indicated rescue constructs. MAP2 labels dendrites. Scale bars= 30 μm. **b,** Quantification of dendritic eIF4G2 immunofluorescence intensity confirms efficient depletion following knockdown and differential restoration of dendritic eIF4G2 signal by the dendritic and somatic rescue constructs used in Fig. 3g,h (n= 3 biological replicates). Data are presented as mean ± SD.

**Extended Data Fig. 8.**
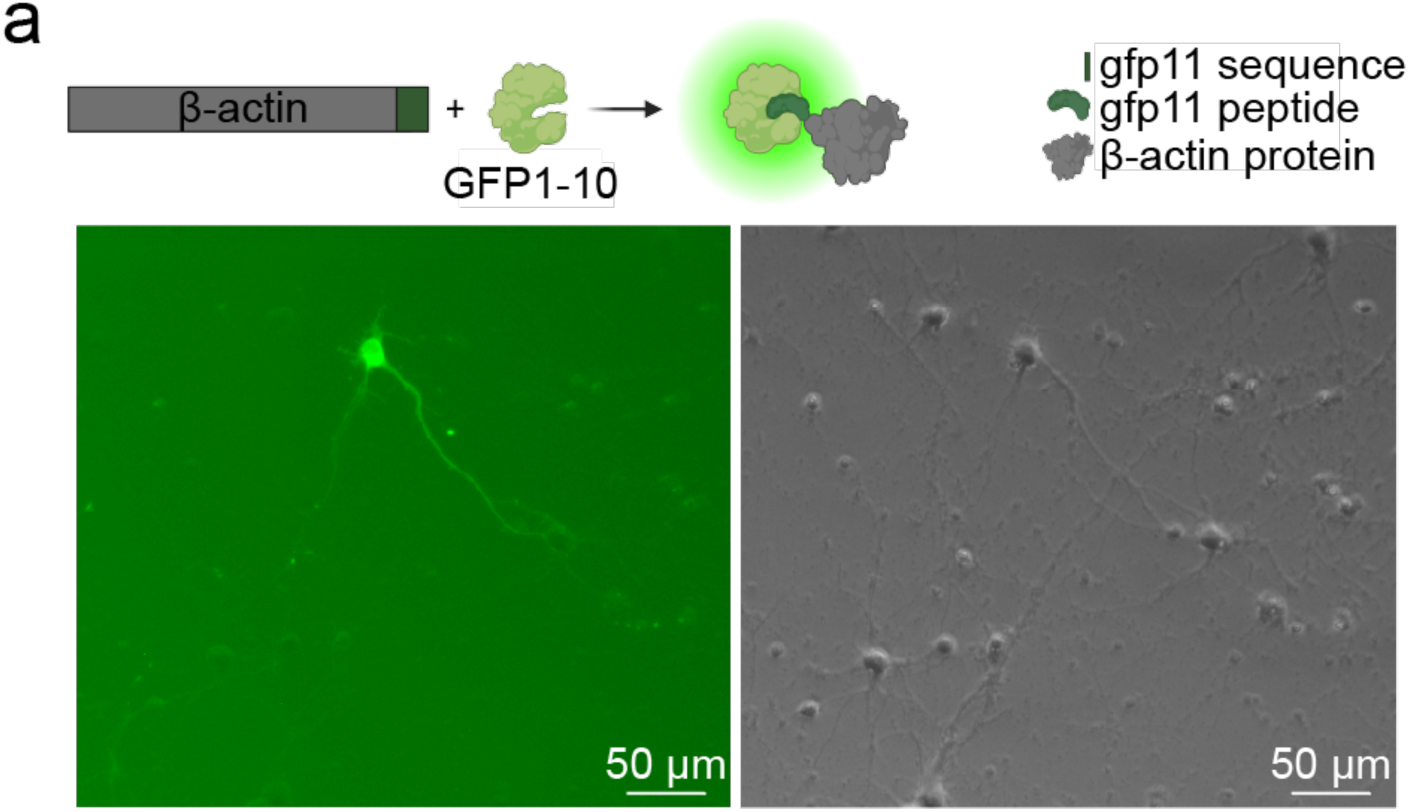
β-actin-GFP11 reporter validates split-GFP complementation. Validation of the GFP1-10/GFP11 complementation system using the β-actin open reading frame fused to a C-terminal GFP11 tag and co-expressed with GFP1-10. Robust GFP fluorescence confirms efficient complementation of the split-GFP system used to detect GFP11-tagged peptide reporters. Scale bars= 50 μm.

**Extended Data Fig. 9.**
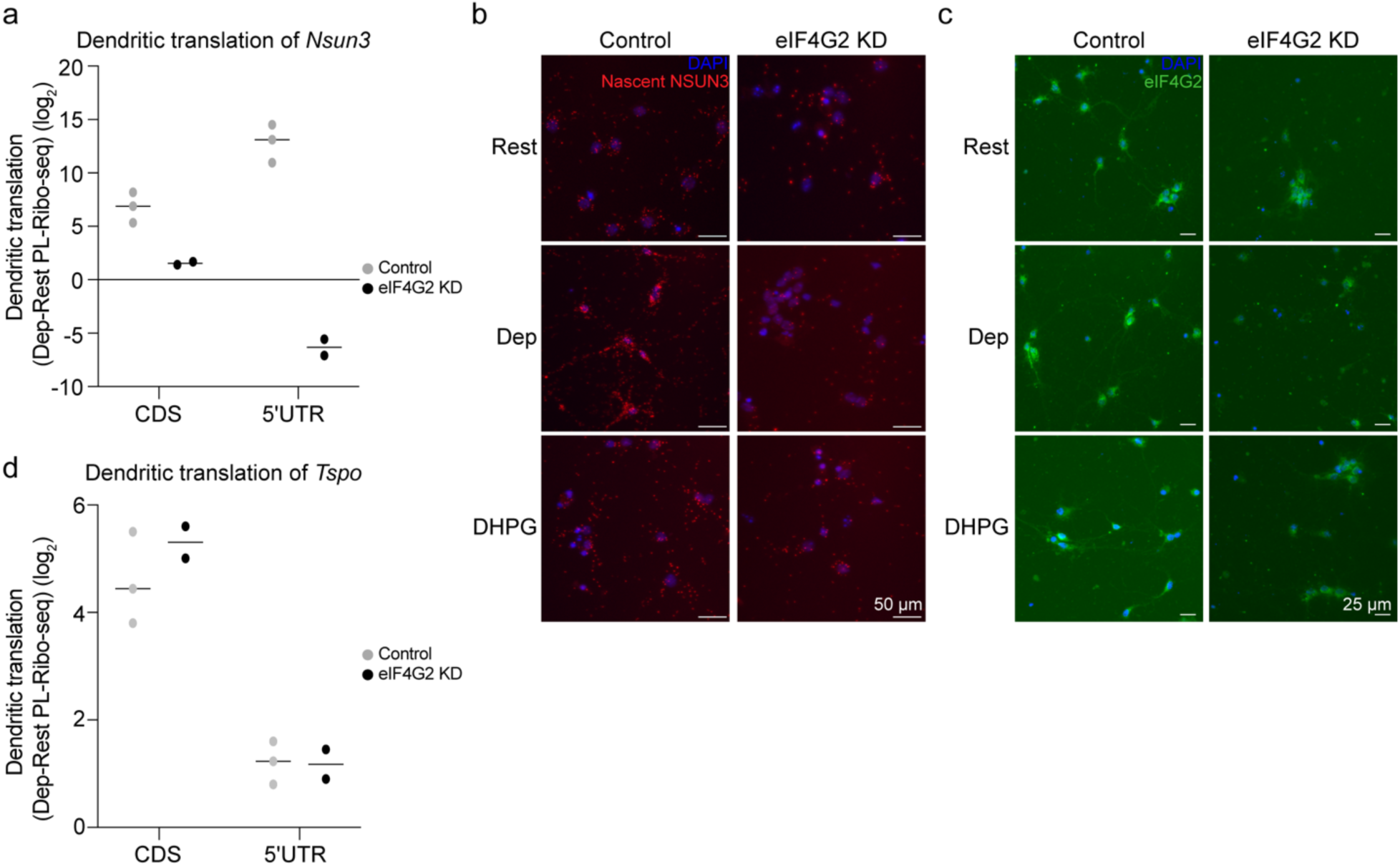
eIF4G2 is required for activity-induced endogenous *Nsun3* translation. **a**, Dendritic translation of endogenous *Nsun3* measured by PL-Ribo-seq in control and eIF4G2 knockdown neurons, shown as Dep-Rest log₂ fold change across the *Nsun3* CDS and 5′ UTR. **b**, Representative puro-PLA images showing activity-induced nascent NSUN3 translation in control neurons and loss of this response following eIF4G2 knockdown under KCl-depolarized and DHPG-treated conditions. Scale bars= 50 μm. **c**, Representative IF images confirming eIF4G2 depletion in knockdown neurons across the corresponding conditions. DAPI labels nuclei. Scale bars= 25 μm. **d**, Dendritic translation of endogenous *Tspo* measured by PL-Ribo-seq in control and eIF4G2 knockdown neurons, shown as Dep-Rest log₂ fold change across the *Tspo* CDS and 5′ UTR.

Data S1. Dendritic eIF4G2 RNA targets enriched in activated versus resting PL-CLIP libraries

Data S2. siRNAs used for knockdown of CAMK2A, eIF4G2, NSUN3, and TSPO

Data S3. Primers used for targeted amplification and Sanger sequencing validation of eIF4G2 PL-CLIP libraries

## Materials and Methods

### Plasmid and reporter cloning

Constructs were cloned into the doxycycline-inducible Lenti-X Tet-On 3G expression system (Takara Bio, 631187) under the control of the PTRE3GS promoter using Gibson assembly (NEB, E5510S). TurboID-PSD95, previously developed in our lab, was used for proximity-labeled CLIP (PL-CLIP) to label postsynaptic compartments. Lentivirus encoding TurboID-PSD95 was generated using Lenti-X Packaging Single Shots (Takara Bio, 631275), as described previously^12^. Myristoylation (Met-Gly-Thr-Val-Leu-Ser-Leu-Ser-Pro-Ser-Tyr) and LDLRct sequences were inserted at the 5′ and 3′ ends, respectively, of *Tspo*, *Nsun3* and *eIF4G2* open reading frames amplified from cDNA. These open reading frames were also fused in-frame to an N-terminal 3xFLAG tag sequence from MilliporeSigma. eIF4G2 phosphorylation-site mutation was introduced using the Q5 Site-Directed Mutagenesis Kit (NEB, E0554S). The somatic localization signal was adapted from Mendonsa et al.^23^ and consisted of the Lynx1 3′UTR sequence amplified from cDNA. GFP11 and GFP1-10 sequences were obtained from Kamiyama et al.^53^ The mCherry coding sequence was amplified from Addgene plasmid #179390, and the β-actin open reading frame was amplified from cDNA. All viral constructs were cloned and propagated at 30°C and verified by Sanger sequencing with forward and reverse primers through Genewiz.

### Primary cortical cultures and neuronal stimulation paradigms

Pregnant CD-1 albino mice (Charles River Laboratories, strain 022) were maintained in accordance with Institutional Animal Care and Use Committee (IACUC) guidelines at The Rockefeller University and the Herbert Wertheim Institute for Biomedical Innovation & Technology. Embryonic day 14.5 (E14.5) cortices were dissected in 1x Hank’s Balanced Salt Solution (HBSS), and primary cortical neurons were prepared as described previously^12^. Neurons were cultured for 14 days in vitro (DIV) on poly-ornithine-coated plates, replacing one-third of the culture medium with fresh medium every 3 days. Unless otherwise indicated, neuronal treatments were performed at 14 DIV. For imaging experiments, neurons were transfected at 8 DIV and treated, fixed, or harvested at 12 DIV. For viral transduction, virus (TurboID-PSD95 and Tet) was added at 2 DIV; after 24 h, half of the medium was replaced with fresh medium and half with conditioned medium, both supplemented with doxycycline at a final concentration of 300 ng/ml. For transient transfections, the culture medium was completely replaced 6 h after transfection with fresh medium containing 300 ng/ml doxycycline.

Neuronal activations were performed as described previously^12^ with minor modifications. Briefly, neurons were silenced by treatment with the sodium channel blocker tetrodotoxin (TTX; Fisher Scientific, 501964082) and 100 μM of the NMDA receptor antagonist DL-2-amino-5-phosphopentanoic acid (DL-AP5; Abcam, ab120271) for 2 hours at 37°C and 5% CO₂. Neurons were subsequently stimulated either by KCl depolarization using a solution containing 170 mM KCl, 2 mM CaCl₂, 1 mM MgCl₂, and 10 mM HEPES in Neurobasal medium, added to a final volume corresponding to 33% of the culture medium, or with the group I mGluR agonist DHPG (Fisher Scientific, 080510) for 30 minutes. Following stimulation, conditioned medium was returned to the cultures for an additional 30 minutes unless otherwise noted. For proximity-labeling experiments, biotin was added to a final concentration of 100 μM during this recovery period. For PL-CLIP experiments, neurons received a 2-minute cycloheximide pulse immediately before biotin addition.

To assess calcium dependence, neurons were pretreated with 10 mM EGTA for 10 minutes at 37°C and 5% CO₂ before stimulation. KCl or DHPG was then added after EGTA to determine whether the activity-induced translational response required extracellular calcium.

### Puromycin proximity ligation assay (puro-PLA)

Nascent protein synthesis was detected using puromycin proximity ligation assays (puro-PLA). Neurons were incubated with 2 μM puromycin (Thermo Fisher Scientific, A1113803) for 10 min at 37°C and 5% CO₂. For activity-dependent translation experiments, puromycin was added during the final 10 min of the 30-minute recovery period following KCl or DHPG stimulation. Cells were washed with pre-warmed PBS-MC (1x PBS, pH 7.4, 1 mM MgCl₂, 0.1 mM CaCl₂), fixed in PBS-MC containing 4% paraformaldehyde and 4% sucrose, and permeabilized with 0.5% Triton X-100 in PBS for 15 minutes as described previously^54^. Puro-PLA was performed using anti-puromycin and protein-specific primary antibodies together with the Duolink In Situ PLA kit (MilliporeSigma, DUO92101) according to the manufacturer’s instructions. Images were acquired on ECHO Revolve R4 and KEYENCE BZ-9000E fluorescence microscopes. PLA puncta were quantified in Fiji using the *Threshold* and *Analyze Particles* functions. Dendritic PLA puncta were quantified within MAP2-positive regions after exclusion of the somatic compartment and normalized to the number of DAPI-positive nuclei in each field of view to obtain the average dendritic signal per neuron.

### Immunofluorescence (IF)

Neurons were cultured on poly-L-ornithine-coated 2-or 4-well Lab-Tek II chambered coverglasses (Nunc). Cells were washed twice with PBS, fixed in 4% paraformaldehyde for 10 minutes at room temperature, permeabilized with 0.2% Triton X-100 in PBS for 10-15 minutes on ice, and blocked for 1 hour at room temperature in 3% donkey or goat serum in 1x PBS, selected according to the secondary antibody host species. Primary antibodies diluted in blocking solution were incubated overnight at 4°C, followed by three 5-minute 1x PBS washes and incubation with species-appropriate secondary antibodies for 2 hours at room temperature. Cells were then washed three times with 1x PBS, with DAPI included during the second wash, and stored in 1x PBS at 4°C in the dark until imaging. Images were acquired using an ECHO Revolve R4 fluorescence microscope. Quantification was performed in Fiji using identical threshold settings across experimental conditions. Background-subtracted fluorescence intensity was measured from manually selected dendritic regions of interest (ROIs) within MAP2-positive regions after exclusion of the somatic compartment and normalized to the number of DAPI-positive nuclei in each image to obtain the average dendritic fluorescence per neuron.

### Mitochondrial mass and membrane potential measurements

Mitochondrial mass and membrane potential were assessed in live neurons using MitoTracker Green FM (Thermo Fisher Scientific, M7514) and MitoTracker Deep Red FM (Thermo Fisher Scientific, M22426), respectively. MitoTracker Green FM and MitoTracker Deep Red FM were added during the final 30 minutes of neuronal silencing at final concentrations of 50 nM and 150 nM, respectively. Resting neurons were incubated with the dyes for the same 30-minute period in regular culture medium. Cells were maintained at 37°C and 5% CO₂ throughout dye loading and stimulation. Following 30 minutes of KCl or DHPG stimulation, or the corresponding period under resting conditions, culture medium was replaced with conditioned phenol red-free Neurobasal medium, and mitochondrial responses were imaged live using an ECHO Revolve R4 microscope within 15 minutes. MitoTracker Green FM was imaged using a FITC/GFP filter set (490/516 nm), whereas MitoTracker Deep Red FM was imaged using a Cy5/far-red filter set (644/665 nm). Fluorescence images were acquired using identical imaging settings across all experimental conditions. Transmitted-light images were acquired simultaneously with the fluorescence channels and overlaid during analysis to facilitate accurate identification and tracing of neuronal dendrites. Processes corresponding to dendrites, identified based on their morphology, branching pattern, and continuity with the dendritic arbor in transmitted-light images, were traced in Fiji, while overlapping neurites, ambiguous processes, and all somatic regions were excluded from analysis. Only clearly identifiable dendritic segments that could be continuously followed throughout the imaging field were included for quantification. Traced dendrites were skeletonized using the Fiji *Skeletonize* function to generate a one-pixel-wide representation of each dendrite, enabling fluorescence measurements to be normalized to the dendritic path length. Mitochondrial fluorescence was quantified exclusively within these dendritic regions and normalized to the total dendritic skeleton length. Normalization to dendritic length was used instead of cell number because fields of view frequently differed in total dendritic coverage, including neurons whose somata were present while portions of their dendritic arbors extended outside the imaging field, as well as images containing variable dendritic densities. This approach minimized technical variability arising from differences in dendritic content between images and provided a stringent, biologically relevant measure of mitochondrial signal per unit dendrite.

### Fluo-4 AM fluorescence measurement

Intracellular calcium levels were assessed in live neurons using Fluo-4 AM (Thermo Fisher Scientific, F14201). Fluo-4 AM was resuspended in DMSO to a stock concentration of 1 mM, and Pluronic F-127 (Thermo Fisher Scientific, P6866) was added at a final concentration of 0.02% to facilitate dye dispersion. Resting and stimulated neurons were first loaded with 2 µM Fluo-4 AM for 45 minutes at 37°C and 5% CO₂. Cells were then washed three times with Neurobasal medium and allowed to de-esterify in dye-free Neurobasal medium for an additional 30 minutes at 37°C and 5% CO₂. Following de-esterification, neurons were either maintained under resting conditions or stimulated with KCl or DHPG for 30 minutes. Fluorescence was measured at the end of this stimulation period while the indicated treatments remained present, thereby capturing sustained intracellular calcium levels during ongoing neuronal activation rather than the initial calcium peak. Fluorescence was measured using a Varioskan Lux plate reader and SkanIt Re 5.0 software at excitation and emission wavelengths of 494 and 506 nm, respectively. Fluorescence from each well was normalized to the corresponding cell number to obtain fluorescence per cell. Each biological replicate represented the mean of three technical replicate wells plated from the same neuronal preparation. Live-cell fluorescence images were also acquired using an ECHO Revolve R4 microscope equipped with a FITC/GFP filter set (490/516 nm), with identical acquisition settings maintained across all conditions within each experiment. For microscopy-based measurements, Fluo-4 fluorescence was quantified on a per-cell basis in Fiji. Individual neuronal cell bodies were outlined as regions of interest using the corresponding transmitted-light images, and overlapping cells or cells with poorly defined boundaries were excluded. Mean fluorescence intensity was measured for each neuron and used as an indicator of sustained stimulus-evoked intracellular calcium levels. Thus, fluorescence measurements obtained using both plate-reader and microscopy-based approaches were calculated on a per-cell basis.

### Reporter transfections and siRNA knockdowns

Primary cortical neurons were transfected at 8 days DIV using Lipofectamine LTX with PLUS Reagent (Thermo Fisher Scientific, 15338030). For each well of a 12-well plate, 0.6 μg of the Tet transactivator construct and 0.6 μg of the reporter construct were co-transfected using 1.6 μl Lipofectamine LTX and 1.2 μl PLUS Reagent. After 4-6 hours, the transfection medium was replaced with fresh culture medium containing doxycycline (300 ng/ml), and cells were analyzed or harvested within 4 days.

For siRNA-mediated knockdown, neurons at 8 DIV and >85% confluency were transfected using Lipofectamine LTX with PLUS Reagent and maintained for 48-72 hours before analysis. siRNAs were used at a final concentration of 20-25 nM, with a non-targeting siRNA included as a control under both resting and activated conditions. For rescue experiments, neurons were first transfected with 15 nM siRNA using 1.4 μl Lipofectamine LTX and 1 μl PLUS Reagent per well to minimize toxicity during the sequential transfection protocol. After 48 hours, neurons were transfected with the indicated dendritically targeted rescue constructs. siRNA sequences are provided in Supplementary Table 2.

### Western blots

Protein lysates were mixed with NuPAGE LDS Sample Buffer (Thermo Fisher Scientific, NP0007) and NuPAGE Sample Reducing Agent (Thermo Fisher Scientific, NP0009), heated at 95°C for 5 min, and resolved on 4-12% Bis-Tris polyacrylamide gels. Proteins were transferred to nitrocellulose membranes using the iBlot 3 Western Blot Transfer System (Thermo Fisher Scientific). Membranes were blocked for 1 hour at room temperature in Intercept TBS Blocking Buffer (LI-COR Biosciences, 927-60001). Primary antibodies were diluted in blocking buffer incubated with membranes overnight at 4°C with gentle agitation. Membranes were washed three times for 5 minutes each in TBST (TBS containing 0.1% Tween-20) and incubated with the appropriate secondary antibodies for 2 hours at room temperature. Membranes were then washed three times in TBST, followed by two additional PBS washes before imaging.

### eIF4G2 proximity-labeled crosslinking immunoprecipitation (eIF4F2 PL-CLIP)

Neurons expressing TurboID-PSD95 were washed twice with 1x PBS containing 100 μg/ml cycloheximide and UV crosslinked on ice in the same buffer using sequential exposures of 400 and 200 mJ/cm^2^. Cells were immediately scraped into ice-cold 1x PBS supplemented with cycloheximide, pelleted by centrifugation (800xg, 10 minutes, 4°C), and either flash-frozen or processed immediately as described previously^12,17,55^ with modifications. Briefly, five, six, and three biological replicates were analyzed, each prepared from six-seven 15-cm culture dishes, for resting, KCl-depolarized, and DHPG-activated neurons, respectively. Cell pellets were lysed in 1 ml PXL buffer (1x PBS containing 0.1% SDS, 0.5% sodium deoxycholate, 0.5% NP-40, and freshly added protease inhibitors). Following DNase treatment as described in Moore et al.^56^, samples were treated with RNase A (Thermo Fisher Scientific, EN0531) at a 1:100 dilution for overdigestion (OD) and a 1:22,500 dilution for underdigestion (UD). eIF4G2–RNA complexes were subsequently immunoprecipitated overnight at 4°C on Protein A Dynabeads (Thermo Fisher Scientific, 10001D) using 12 μg of rabbit polyclonal anti-eIF4G2 antibody (Thermo Fisher Scientific, PA5-21377) per sample. The bead-bound complexes were subjected to stringent washes as previously described^55^. Following RNA linker ligation and ^32^P-γ-ATP labeling, the Protein A beads were washed twice with PXL buffer and twice with 5x PXL buffer (5x PBS, 0.1% SDS, 0.5% sodium deoxycholate, 0.5% NP-40) before elution in 1% SDS/PXL buffer for 10 minutes at 70°C with shaking at 1100 rpm. Eluates were diluted tenfold in 1x PBS and incubated overnight at 4°C with 75 μl of prewashed Dynabeads MyOne Streptavidin C1 magnetic beads per sample (Thermo Fisher Scientific, 65001) in the presence of 20 U/ml SUPERase-In RNase inhibitor (Thermo Fisher Scientific, AM2694). Streptavidin beads were washed twice with PXL buffer and twice with PNK-EGTA buffer (40 mM Tris pH 7.0, 10 mM Tris pH 8.0, 20 mM EGTA, 0.5% NP-40). Biotinylated complexes were eluted by competition with 2 mM biotin in PNK-EGTA buffer containing NuPAGE LDS Sample Buffer and NuPAGE Sample Reducing Agent. SDS-PAGE, nitrocellulose transfer, and RNA recovery were performed as described previously^56^. cDNA synthesis, circularization, and PCR amplification were carried out as described previously^12^. Following reverse transcription and barcode incorporation, resting and activated samples from each biological replicate were pooled to maximize library yield. Libraries were sequenced on an Illumina NextSeq 2000 using a P1 flow cell.

### Targeted amplification and validation of CLIP libraries

For targeted validation of CLIP libraries, 1-2 µl of each library, at approximately 200-300 pg/µl, was used as template for PCR amplification with gene-specific primers listed in Supplementary Table 3. Reactions were amplified for 30 cycles using the corresponding forward and reverse primer pairs. PCR products were resolved on 1% agarose gels, and DNA fragments within the expected 100-300-bp size range were excised from regions containing a discrete amplification product. Gel-purified products were submitted to Genewiz for Sanger sequencing. Resulting sequences were aligned to the mouse genome using BLAT to confirm the identity and genomic origin of the amplified CLIP products.

### Oxygen consumption measurement

Neuronal oxygen consumption was measured using the Oxygen Consumption Rate Assay Kit (Cayman Chemical, 600800) according to the manufacturer’s instructions. Primary cortical neurons were plated in 96-well plates with six technical replicates per condition and three independent biological replicates. Resting cultures were maintained in conditioned medium without silencing or stimulation. For activated conditions, neurons were silenced as described above and subsequently treated with KCl or DHPG, which remained present throughout the assay. The phosphorescent oxygen probe and assay reagents were then added according to the manufacturer’s protocol, and oxygen consumption was monitored for 120 minutes with measurements acquired every 15 minutes using a Varioskan LUX multimode microplate reader (Thermo Fisher Scientific). Data acquisition and analysis were performed using SkanIt RE 5.0 software. Oxygen consumption rates were determined from the time-dependent increase in probe phosphorescence, which is inversely proportional to extracellular oxygen concentration.

### Antibodies

Primary antibodies: Puromycin (1:3000, mouse, Kerafast, EQ0001, RRID: AB_2620162), Flag (1:3000, mouse, Sigma, F1804, RRID: AB_262044), β-Actin (1:2500, mouse, Sigma, A1978, RRID:AB_476692), MAP2 (1:2500, guinea pig, Synaptic Systems, 188004, RRID: AB_2138181), eIF4G2 (for IF: 1:250, rabbit, Cell Signaling, 5169, RRID: AB_10622189; for PL-CLIP: rabbit, Thermo Fisher Scientific, PA5-21377, RRID: AB_11154244), TSPO (1:200 (PLA), 1:500 (Western blots), Thermo Fisher Scientific, MA5-24844, RRID: AB_2717282, NSUN3 (1:200 (PLA), 1:500 (Western blots), rabbit, LSBio, LS-C747965), SLC25A31 (1:250, biorbyt, orb670517), MFF (1:500, Cell Signaling, 84580, RRID:AB_2728769).

Donkey anti-rabbit 800 (LI-COR Biosciences, 926-32213), donkey anti-rabbit 680 (LI-COR Biosciences, 926-68073), goat anti-mouse 800 (LI-COR, 926-32210), goat anti-mouse 680 (LI-COR Biosciences, 926-68070), donkey anti-mouse Alexa Fluor 488 (Thermo Fisher Scientific, R37114, RRID: AB_2556542), donkey anti-guinea pig Alexa Fluor 488 (Jackson ImmunoResearch Labs, 706-545-148, RRID: AB_2340472), donkey anti-rabbit Alexa Fluor 647 (Jackson ImmunoResearch Labs, 711-605-152, RRID: AB_2492288), donkey anti-mouse Alexa Fluor 555 (Thermo Fisher Scientific, A-31570, RRID: AB_2536180).

### Bioinformatics

#### Principal component analysis

Principal component analysis (PCA) was performed in R using the *prcomp* function on the 500 most variable genes across samples. The number and relative contribution of principal components were evaluated using the pcascree function. PCA plots were generated with the *fviz_pca_ind* function from the factoextra package.

#### Gene set enrichment analysis (GSEA)

Gene set enrichment analysis (GSEA) was performed in R using the *fgsea* package from Bioconductor to identify biological pathways and functional gene sets enriched across the indicated dendritic datasets. Genes were ranked according to the relevant differential-analysis statistic for each comparison, and enrichment was assessed using the corresponding ranked gene lists. Unless otherwise indicated, analyses were performed using the default *fgsea* parameters. Gene sets with a Benjamini-Hochberg false discovery rate (FDR) below 0.05 were considered significantly enriched. The gene-set collections and ranking metrics used for each analysis are specified in the corresponding figure legends.

#### PL-CLIP sequencing and analysis

PL-CLIP libraries were sequenced on an Illumina NextSeq 2000 to generate 75-nt single-end reads. Reads were processed as described previously^56,57^, using the CTK software package for quality filtering, demultiplexing, removal of 5′ and 3′ linker sequences, and collapsing of exact duplicates. Processed reads were aligned to the mouse genome (mm10) using the *align* function from the Rsubread package^58^, allowing up to five mismatches and requiring a minimum aligned read length of 20 nucleotides. For peak-level analyses, reproducible CLIP peaks identified by the CTK pipeline were imported into R using the *rtracklayer* package and annotated to genomic features, including 5′ untranslated regions (5′UTRs), coding sequences (CDSs), 3′UTRs, introns, and intergenic regions. For each biological replicate, the fraction of annotated peaks mapping to each genomic feature was calculated and normalized to the total number of peaks within that library. Replicate-normalized peak distributions were then averaged across biological replicates to generate the genomic peak-density profiles shown in the figures. To quantify activity-dependent eIF4G2 binding at the transcript level, annotated 5′UTR regions were obtained from the *TxDb.Mmusculus.UCSC.mm10.knownGene* package, and overlapping eIF4G2 PL-CLIP reads were quantified using *summarizeOverlaps* from the *GenomicAlignments* package^59^. For genes with multiple annotated isoforms, the longest transcript was used. The number of reads mapping to each 5’UTR was normalized for library depth, and log_2_ (fold change, KCl vs. Rest and DHPG vs. Rest) values were calculated using a pseudocount of 0.1. Statistical significance of differential CLIP-tags between conditions was assessed using binomial tests based on the normalized CLIP-tag counts.

